# Direct interoceptive input to the insular cortex shapes learned feeding behavior

**DOI:** 10.1101/2025.05.13.653896

**Authors:** Zhe Zhao, Binbin Xu, Skylar Anthony, Suganya Subramanian, Bryan Granger, Carolyn Von-Walter, Elisa Mizrachi, Dhruvum Bajpai, Paul Tyagi, Matthew Kidd, Abhishikta Srigiriraju, Isaac McKie, Zhiying Li, M. McLean Bolton, Stefano Berto, Sarah A. Stern

**Affiliations:** Max Planck Florida Institute for Neuroscience, Jupiter, FL 33458, USA; EuroMov Digital Health in Motion, Univ Montpellier, IMT Mines Alès, 6 Av. de Clavières, 30100 Alès, France; Medical University of South Carolina, Charleston, SC 29425, USA; International Max Planck Research School for Synapses and Circuits, Jupiter, FL 33458, USA; University of Florida, Gainesville, FL 32611, USA; Florida Atlantic University, Boca Raton, FL 33431, USA; The Rockefeller University, New York, NY 10065, USA

**Keywords:** insular cortex, leptin, feeding, body weight, operant behavior, interoception, calcium imaging, transcriptomics, optogenetics

## Abstract

The insular cortex (insula) is an interoceptive hub, which senses internal states such as hunger, thirst, pain, and emotions. Previous studies suggest that the insula directly senses internal states, but the mechanisms remain elusive. We identified a population of leptin receptor-positive cells with a unique morphology in the insula (INS^LepR^). Based on leptin’s known role in signaling adiposity, we hypothesized that INS^LepR^ neurons detect internal states to regulate food intake and body weight. Accordingly, we found that intra-insula leptin administration or optogenetic stimulation of INS^LepR^ neurons impacts feeding behavior. Moreover, INS^LepR^ neuron activity encodes feeding bouts in an internal-state dependent manner, and leptin alters insula neural dynamics in response to feeding, while also reshaping the transcriptome. Taken together, our data supports a model for direct interoceptive input to the insula, in which INS^LepR^ cells integrate adiposity level signals to regulate feeding and body weight in a learned manner.

## INTRODUCTION

Food intake is a complex process involving both interoceptive detection of internal hunger states, but also exteroceptive detection of sensory cues, potential predators, social stimuli and others. The insular cortex (insula) is a multisensory brain region, known to be involved in diverse functions such as taste, hunger, thirst, pain, anxiety and social behavior^1–6^. It is hypothesized to play a role in both the exteroceptive parts of behavior, in the form of salience detection^7–9^, as well as the interoceptive aspects such as detection of bodily states^10–13^. The role of the insula in food intake is complex; this brain region, also referred to as the gustatory cortex, has primarily been studied because it contains neurons which encode taste^14–18^. However, numerous studies have shown that the insula is not required for homeostatic food intake, but is required for complex, learned ingestive behavior such as conditioned taste aversion, conditioned overconsumption and palatable food intake^19–22^. Nevertheless, studies have reported that activation of specific projections from the insula may lead to decreased food intake^23,24^, although there is some concern regarding the non-physiological nature of the optogenetic activation paradigms used. One longstanding hypothesis is that the insula is an interoceptive hub, and that its role is to integrate the information about internal state with information about the external world to drive complex behaviors.

Indeed, one recent study showed that neural activity in the insula is correlated with thirst and satiety states, but this activity is unchanged by activation of subfornical organ thirst neurons^25^, suggesting that the insula can directly track interoceptive state. However, the mechanisms driving these responses are unknown. Previous studies suggest that internal state information arrives at the insula indirectly through vagus nerve projections^26–28^. However, a direct mechanism for internal state sensing has not been ruled out. Here we identify leptin receptors (LepR) in the insula as such a candidate.

Leptin receptors are known primarily for their role in the hypothalamus, sensing leptin, which is released from adipose tissue and reflects total amount of body fat^29–31^. It has long been known that there are moderate amounts of leptin receptors in areas outside of the hypothalamus, such as the hippocampus and ventral tegmental area^32–34^, but the role of extrahypothalamic LepR remains elusive. Here we find that LepR marks a unique subset of neurons mainly located in the deep layers of the insula and claustrum (INS^LepR^), and that the interoceptive input provided by leptin influences food intake and body weight, primarily to coordinate learned feeding behavior. Injection of leptin directly into the insula decreases food intake and body weight by increasing the intrinsic excitability of INS^LepR^ cells. Using both bulk and single-cell transcriptomics, we found that LepR is expressed in a unique subset of recently identified glutamatergic Car3 (carbonic anhydrase III)-expressing neurons located primarily in deep layers bordering the claustrum region, as well as on vascular cells that mediate active transport across the blood-brain barrier. Using *in vivo* calcium imaging, we demonstrated that activity of INS^LepR^ cells reliably decoded feeding behavior, and INS^LepR^ neurons are tuned to various aspects of both food consumption and operant nose-poking for food. Activation of INS^LepR^ cells altered food intake in an operant food reward paradigm, without affecting homeostatic feeding. Transcriptomic analysis revealed that systemic leptin treatment led to numerous changes in metabolic and feeding-related pathways in the insula and reshaped the population activity of the insula during food consumption. Lastly, we found that INS^LepR^ cells project within the insula and to the basolateral amygdala, another area with a known role in food intake and reward^35,36^. Thus, leptin receptors emerge as interoceptive detectors in the insula for internal states, which enables learned feeding behaviors.

## RESULTS

### Leptin infusion to the insula regulates food intake and body weight

We first examined LepR expression in the insula using a LepR-Cre mouse crossed with a TdTomato reporter line, and observed a significant expression of TdTomato exclusively the insula and claustrum area, but not other cortical areas (Figure S1A). We confirmed this finding using fluorescent in situ hybridization (Figure 1A), which showed significant LepR expression in the layer 5/6 region of the insula as well as in the claustrum. We further examined the neurotransmitter identity of the insula-claustrum LepR+ neurons (INS^LepR^) and found that they were almost exclusively glutamatergic (Figures 1A and S1B-E). We then quantified the number of LepR+ cells across layers following an injection of cre-dependent mCherry virus into the insula of LepR-Cre mice and registration of mCherry+ cells to Nissl stained cell bodies (Figures 1B and S1F). We confirmed that the majority of mCherry+ cells were in deeper layers of the insula, with some sparse expression in the superficial layers (Figures 1C-D, Video S1).

**Figure 1.**
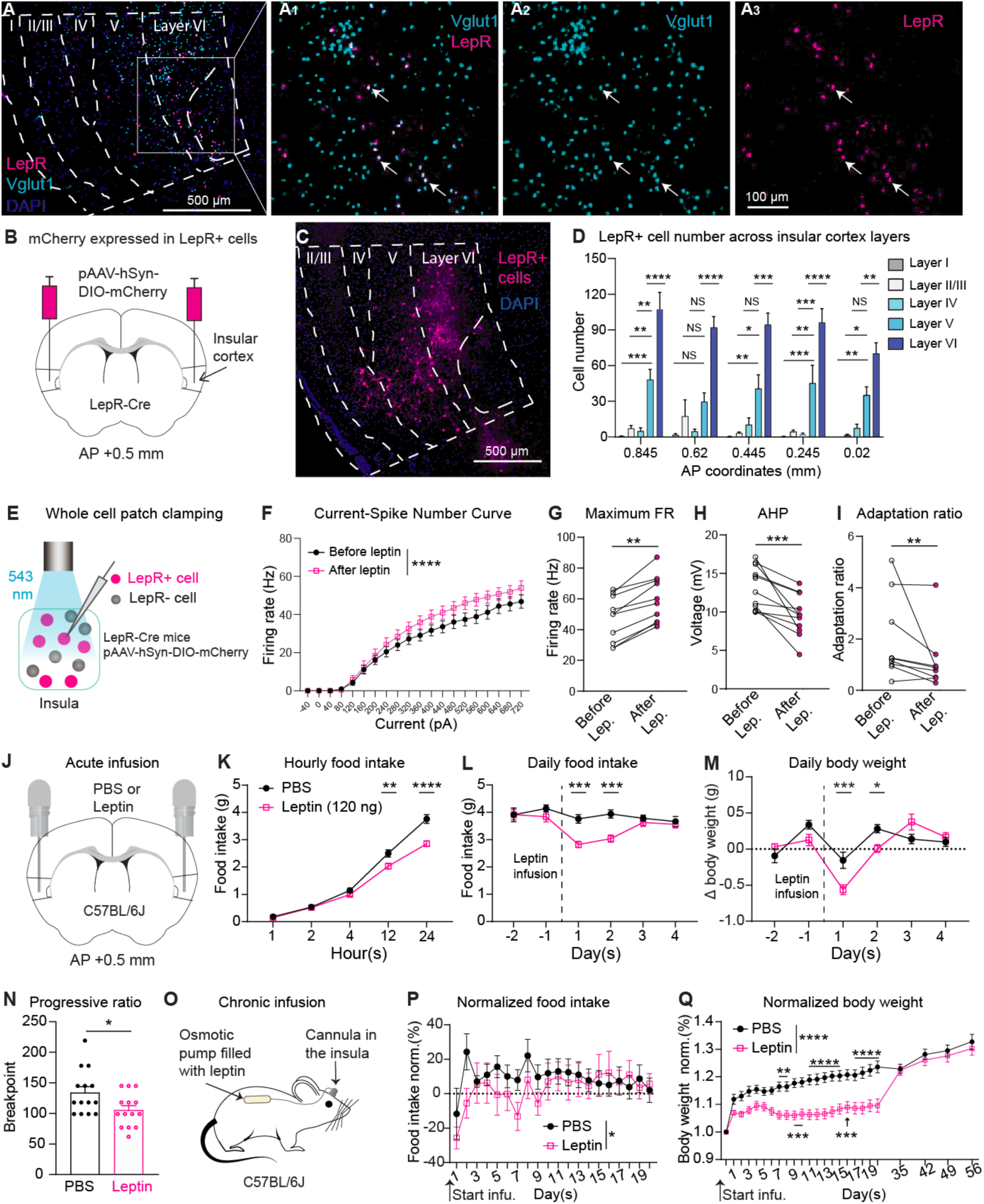
Leptin increases the excitability of INS^LepR^ neurons and modulates food intake, body weight and motivation. (A) Two-color fluorescent *in situ* hybridization for LepR mRNA (magenta) and Vglut1 mRNA (cyan) in the insula-claustrum region. A_1-3_ (inset from A) show LepR mRNA and Vglut1 mRNA in deep layers of the insula. White arrows point to colocalized neurons. (B) Scheme for Cre-dependent mCherry virus injected bilaterally into the insula of LepR-Cre mice. (C) mCherry (magenta) expression in LepR+ cells located the insula. (D) LepR+ cell number across the anterior/posterior (AP) axis of the insular cortex (n = 6-8/group, two-way ANOVA followed by Turkey post-hoc test, F_(4,160)_ = 125.6, **** *p* < 0.0001. Layer I vs. Layer6 VI, Layer II/III vs. Layer6 VI, Layer IV vs. Layer VI at all coordinates in the graph, **** *p* < 0.0001. Layer I vs. Layer V, Layer II/III vs. Layer V, Layer IV vs. Layer V, Layer V vs. Layer VI at different coordinates, \**p* < 0.05, \*\**p* < 0.01, *** *p* < 0.001, **** *p* < 0.0001, NS: No significance). (E) Scheme for whole cell patch-clamp recording of LepR+ cells. (F) Firing rate (expressed in hertz, Hz) triggered by a series of currents before and after leptin administration (n = 7-10 cells from 7 mice in each group, two-way ANOVA, F_(1,345)_ = 25.7, **** *p* < 0.0001). (G) Maximum firing rate (expressed in hertz, Hz) triggered by current injection before and after leptin (Lep.) administration (n = 10 from 7 mice, Wilcoxon matched-pairs signed rank test, ** *p* = 0.002). (H) Afterhyperpolarization (AHP) in the first action potential triggered by the minimum current, before and after leptin (Lep.) administration (n = 11, Wilcoxon matched-pairs signed rank test, *** *p* = 0.001). (I) Adaptation ratio (inter spike interval (ISI) of the 5^th^ and 6^th^ spikes/ISI of 1^st^ and 2^nd^ spikes) before and after leptin (Lep.) administration (n = 9, Wilcoxon matched-pairs signed rank test, * *p* = 0.0117; excluded one outlier). (J) Scheme for cannula implantation and local infusion of PBS or leptin (120ng) into the insula. (K) Food intake measurements, expressed in cumulative grams (s) consumed following infusion of PBS or leptin (120ng) into the insula (n = 16 DPBS, n = 16 leptin, two-way ANOVA followed by Turkey post-hoc test, F_(1,150)_ = 31.06, **** *p* < 0.0001. PBS vs Leptin, at 12-hour, ** *p* = 0.0014; PBS vs Leptin, at 24-hour, **** *p* < 0.0001). (L-M) Food intake, expressed as grams (g) per day, and body weight measurements, expressed as the grams (g) change (Δ) in body weight from baseline (-2 days) before and after the injection of PBS or leptin (120ng) into the insula-claustrum region (For **L**, n = 15-16 PBS, n = 16 leptin, two-way ANOVA, PBS vs. leptin, F_(1,179)_ = 18.71, **** *p* < 0.0001; at 1^st^, *** *p* = 0.0003; at 2^nd^, *** *p* = 0.0006. For **M**, n = 16 PBS, n = 16 leptin (Days -2 to 2, 4) n = 8 PBS, n = 8 leptin (Day 3); DPBS vs. leptin, two-way ANOVA, F_(1,164)_ = 3.025, *p* = 0.0838; at 1^st^, *** *p* < 0.0005; at 2^nd^, * *p* = 0.0407). (N) Breakpoint, represented as the maximum # of nose pokes required to get a food pellet in the progressive ratio test, 1 day after PBS or leptin infusion (n = 14 PBS, n = 14 leptin, two-tailed unpaired t test, * *p* = 0.027). (O) Scheme for chronic local infusion of PBS or leptin (5ng/hour for ∼14 days) using osmotic pumps. (P-Q) Food intake and body weight, normalized to the average daily food intake and body weight in the week prior to osmotic pump implantation, expressed as the % increase in grams (g) from starting body weight, after chronic PBS and leptin infusion to the insula (For **P**, n = 15-16 PBS, n = 15-16 leptin, two-way ANOVA followed by Turkey post-hoc test, PBS vs. leptin, F_(1,356)_ = 5.54, * *p* = 0.0191. For **Q**, n = 15-16 PBS, n = 15-16 leptin, PBS vs. leptin, two-way ANOVA, F_(1,450)_ = 290.6, **** *p* < 0.0001. *** *p* < 0.001, \*\**p* < 0.01). For more information, see **Figures S1** and **S2**.

Although previous findings also concluded that there is sparse leptin receptor expression in the cortex^32–34^, they have been largely unexplored as an important factor in regulating food intake and body weight^37^. One previous study demonstrated that leptin administration enhances synaptic transmission in the rat insula, but they did not examine whether INS^LepR^ cells mediated this effect^38^. Another study reported that leptin shapes the intrinsic excitability of pyramidal neuron in rat insular cortex^39^, but it is unknown how leptin affects INS^LepR^ cells’ electrophysiological properties, nor the behavioral consequences of leptin administration. We first asked whether leptin changes intrinsic neural excitability in the insula. We applied slice electrophysiology using whole cell patch clamp (Figures 1E and S2A) and found that superfusion of leptin increases the intrinsic excitability of INS^LepR^ cells, with a significant increase in the firing rate induced by current injection, maximum firing rate, decrease in after hyperpolarization and adaptation ratio (Figures 1F-I). Conversely, leptin administration decreased intrinsic excitability in leptin receptor-negative (LepR-) cells (Figures S2B and C), confirming that leptin indeed alters INS^LepR^ neural excitability in the insula.

We next asked whether direct bilateral infusion of leptin into the insula would impact food intake and body weight (Figures 1J and S2D). We assessed food intake using the automated feeding dispenser FED3.0^40^, which enabled us to have precise quantification of food intake during experiments. We found that a single acute infusion of leptin led to a decrease in food intake that began around 12 hours after injection (Figure 1K) and lasted for two days (Figure 1L), before returning to baseline. This was accompanied by a small, but significant decrease in body weight as well (Figure 1M). In addition, we also used the FED3.0 to assess motivation. A single acute infusion of leptin led to a decrease in breakpoint in a progressive ratio nose-poke task, indicating that mice had less motivation to work for food pellets when compared to controls (Figures 1N, S2E and F). This effect also lasted 2 days (Figure S2E) and was normalized by 6 days after the infusion (Figure S2F). In addition, because endogenous leptin is released continuously in the bloodstream^41^, we implanted osmotic pumps with a cannula directly into the insula to infuse leptin continuously over 14 days (Figure 1O). Chronic infusion of leptin led to a small reduction in food intake, particularly in the first 9 days (Figure 1P), and a significant decrease in body weight through the entire infusion period, relative to controls (Figure 1Q). Following removal of the pump, body weight normalized and was again equivalent to PBS-injected controls by 35 days following the surgery. In addition, the effect of leptin was dependent on the mouse’s internal state at the time of infusion; infusion after an 18-hour food restriction led to a significant decrease in food intake between 4 and 48 hours after the infusion (Figure S2G), which was not observed when leptin was infused before an 18-hour food restriction (Figure S2H).

Activation of INS^LepR^ neurons disrupts learned feeding and induces avoidance We next examined the consequences of manipulating INS^LepR^ neurons directly. We first chemogenetically activated and inhibited INS^LepR^ neurons, but did not see any changes in homeostatic food intake, measured in the homecage (Figures S3A-B). We then ablated INS^LepR^ neurons by injecting cre-dependent Caspase3 (Casp3) into the insula of LepR-Cre mice (Figure S3C). Although we observed successful ablation (Figure S3D), there was no difference between Casp3 injected animals and controls in body weight over 11 weeks (Figures S3E), suggesting that INS^LepR^ are not required to regulate overall homeostatic feeding. Because previous studies^19–21^ indicate that the insula may regulate more complex learned feeding behaviors rather than homeostatic food intake, we hypothesized that INS^LepR^ neurons might instead provide interoceptive information that shapes learned feeding behaviors. To examine this, we used a chamber developed by our group, called INGEsT^42^ (Figure 2A), and mice were trained in a task in which they learned to nosepoke followed by food or water availability (Figure 2B). LepR-Cre mice were injected with cre-dependent ChR2 into the insula alongside bilateral implantation of optical fibers (Figure 2C). Mice were then food restricted and trained to nosepoke for food. After the operant feeding test, mice were switched to water restriction for the operant drinking test (Figures 2B and S3F). Mice were subsequently tested with two different stimulation protocols. In the first, we optogenetically stimulated the neurons for 10 minutes prior to the behavioral session (Figure 2D). This led to a decrease in the number of pellets retrieved following a nose-poke, but no change in water intake (Figures 2E and F). We also examined the effect of stimulating LepR+ neurons for 2 seconds after the nose poke before the pellet or lick spout is made available (Figure 2G). This manipulation led to a non-significant increase in the time between nosepoke and pellet retrieval, but no change in the time to water consumption (Figures 2H and I). Thus, activation of LepR+ neurons decreases operant-based feeding, but not drinking. In both cases, stimulation with light in mCherry control mice had no effect on either stimulation protocol (Figures S3G-K).

**Figure 2:**
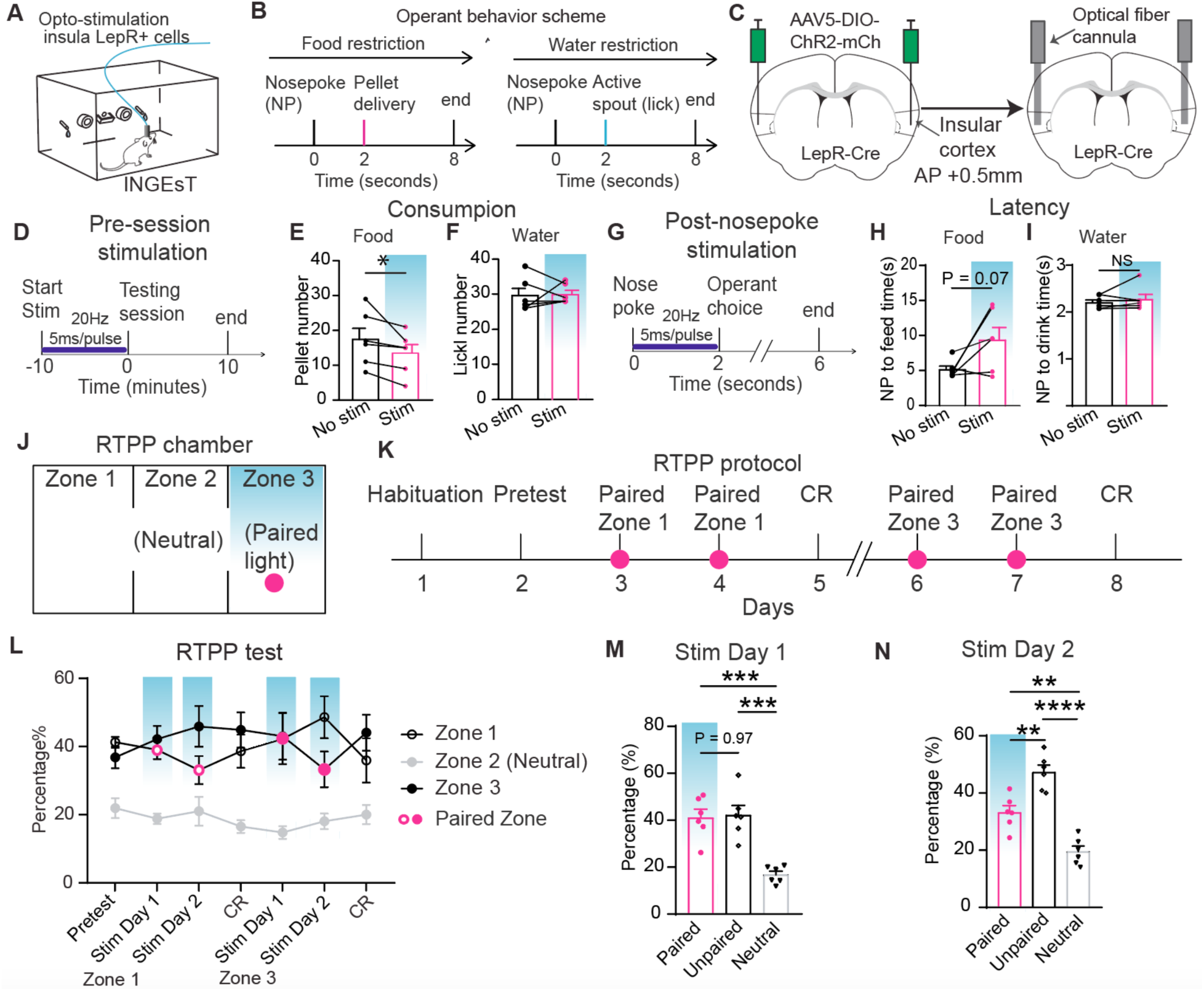
Stimulation of INS^LepR^ cells in the insula suppresses learned operant feeding. (A) Scheme for optogenetic stimulation experiments in the INstument for Gauging Eating and Thirst (INGEsT) chamber. (B) Scheme for operant feeding task. Following the nose poke, a pellet is delivered and the water spout becomes active. Mice may then choose to lick, eat or do both (lick/feed). (C) Scheme for viral injection and optical fiber cannula implantation for optogenetic stimulation of LepR+ cells in the insula. (D) Scheme for pre-session optogenetic stimulation (blue). (E) Pellet number consumed over one testing session with pre-session stimulation (magenta) or control (blue) (n = 6 Stim, n = 6 No stim, two-tailed paired t test, * *p* = 0.0136). (F) Number of licks taken over one testing session with pre-session stimulation (magenta) or control (blue) (n = 6 Stim, n = 6 No stim, two-tailed paired t test, *p* = 0.8682). (G) Scheme for the protocol of post-nosepoke stimulation. Stimulation is given for 2 seconds following the nosepoke before food and water choice is available. (H) Latency to pellet retrieval following nose poke with stimulation (magenta) or control (blue) (n = 6 Stim, n = 6 No stim, two-tailed paired t test, * *p* = 0.0712). (I) Latency to licking water after nose poke with stimulation (magenta) or control (blue) (n = 6 Stim, n = 6 No stim, two-tailed paired t test, * *p* = 0.3917). (J) Scheme for the real-time place preference (RTPP) chamber coupled with optogenetic stimulation. Stimulation is depicted in Zone 3, but switches between Zone 1 (Days 3 and 4) and Zone 3 (Days 6-7). (K) Scheme for the RTPP protocol. The pretest measures initial chamber preference. CR: conditioned response, measures memory for the stimulated side. (L) Percentage of time staying in Zone 1-3 on different days in the RTPP protocol. The stimulated zone is listed under the relevant days on the x-axis. (M) Percentage of time staying in the paired (stimulation) and unpaired (unstimulated) zones on Stim Day 1 (n=6, one-way ANOVA, F_(2,15)_ = 36.75, **** *p* < 0.0001; Paired vs. Unpaired, *p* = 0.9653; Paired vs. Neutral, *** *p* = 0.0003; Unpaired vs. Neutral, *** *p* = 0.0002). (N) Percentage of time staying in Zone 1-3 on Stim Day 2 (n=6, one-way ANOVA, F_(2,15)_ = 19.10, **** *p* < 0.0001; Paired vs. Unpaired, ** *p* = 0.0016; Paired vs. Neutral, ** *p* = 0.002; Unpaired vs. Neutral, **** *p* < 0.0001). For more information, see **Figure S3**.

We then examined whether activation of INS^LepR^ neurons induces approach or avoidance using real time place preference (Figure 2J). Mice were placed in a chamber with 2 compartments and a middle zone, with one compartment paired with stimulation (Figure 2K). Mice were habituated to the chamber and checked for an initial preference, after which INS^LepR^ neurons were stimulated for 2 sessions. They were then tested without stimulation for a conditioned response, after which the stimulation side was reversed and the test repeated (Figure 2K). We found that two days of stimulation induced avoidance of the stimulated side (Figures 2L-N, and S3L-M). Here, too, stimulation in mCherry control mice did not lead to any avoidance (Figures S3N-P). Thus, activation of INS^LepR^ cells induces avoidance in a delayed manner.

Overall, these results suggest that leptin receptors in the insula modulate food intake, particularly when learning is involved.

### INS^LepR^ neural activity encodes operant feeding and is shaped by hunger state

Because we saw the INS^LepR^ neuron activation decreased food reward retrieval in the INGEsT chamber, we wanted to know how these neurons encode task-related information. We conducted calcium imaging of INS^LepR^ neurons using GCaMP8m calcium indicator and a GRIN lens implanted directly into the insula, coupled with a miniscope in freely behaving mice, tested in a similar fixed-ratio operant-based learning task (Figures 3A-C). Mice that were food restricted to 85-90% of their normal body weight were first habituated to the arena and then were trained to nose poke followed by a 2-second delay, after which a chow pellet was delivered from a FED3.0 device.

**Figure 3.**
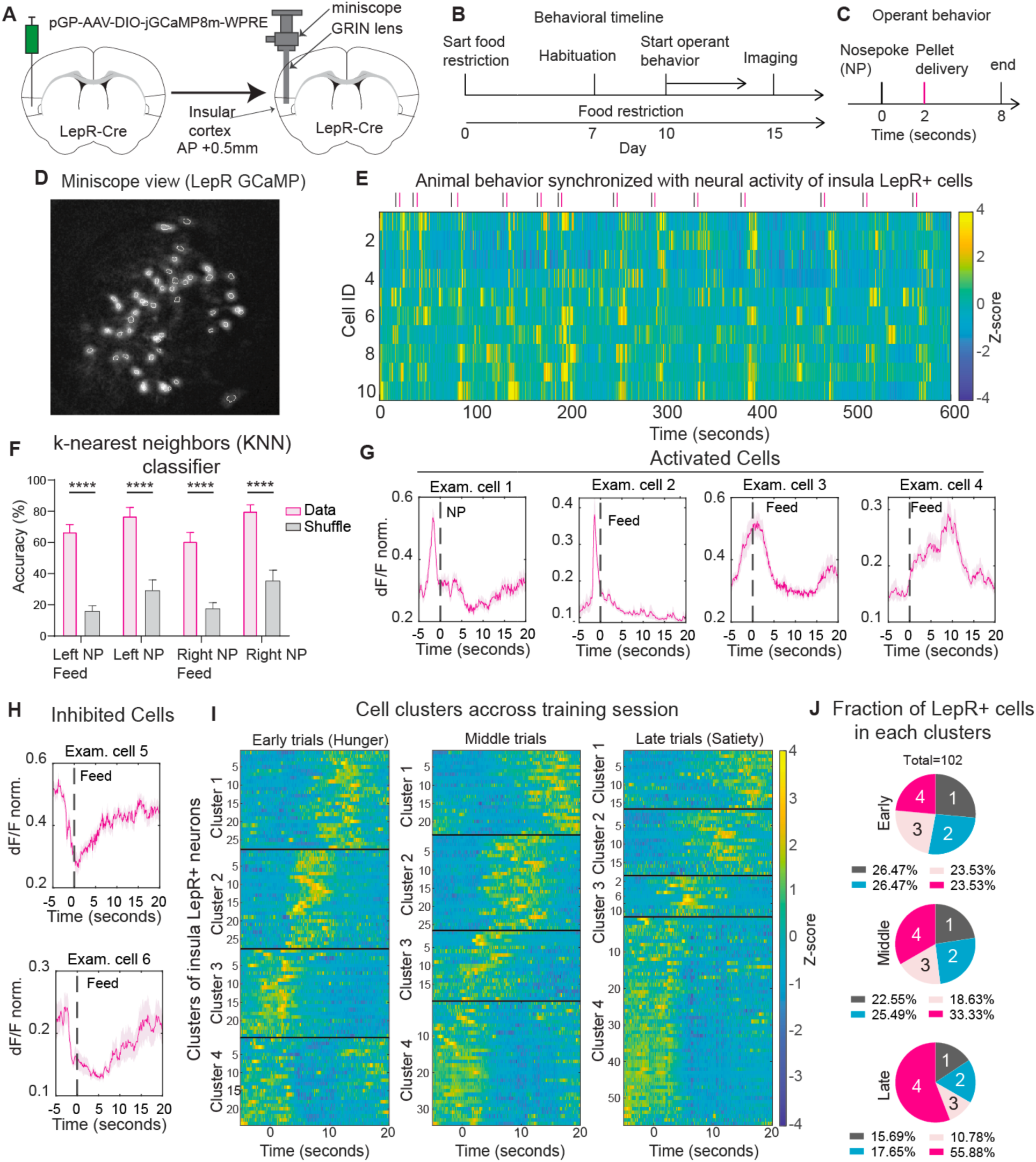
LepR+ cells in the insula encode feeding bouts and track satiety state. (A) Scheme for the method of viral injection and GRIN lens implantation in the insula of LepR-Cre mice. (B) Behavioral training timeline of operant feeding. (C) Scheme for operant feeding task. NP: nose poke. (D) Miniscope view of GCaMP calcium signals expressed in LepR+ cells (maximum projection of fluorescence). (E) Example from one mouse of calcium traces (represented as z-score) from individual cells synchronized to behavior behaviors in one testing session. Behavior is overlayed on top as blue (nosepoke) or pink (pellet retrieval) lines. Each row represents one cell during the entire session length. (F) Percent accuracy of a k-nearest neighbors (KNN) classifier trained by a population of LepR+ cell activity, comparing real data (pink) to shuffled data (gray) (n = 6 data, n = 6 shuffle, data vs. shuffle, two-way ANOVA followed by Turkey post-hoc test, F_(1,40)_ = 160.6, **** *p* < 0.0001; decoding accuracy for Left NP Feed, Left NP, Right NP Feed, and Right NP, **** p< 0.0001). (G-H) Example cells in response to nose poke (NP) or pellet retrieval (Feed) (**G**, activated cells; **H**, inhibited cells). (I) Classification of LepR+ cells aligned to nose poke followed by feeding at early, middle and late trials defined as the first, middle and last third of trials in a testing session, by using hierarchical cluster analysis. Each row represents activity from one cell. (J) Fraction of cells during each phase (early, middle, or late) in each of the four clusters. For more information, see **Figure S4**.

Mice freely choose between a left or right nose-poke port, and reliably retrieved pellets shortly after initiating the nosepoke (Figures S4A-C). Synchronizing neural activity to behavior demonstrated that there were clear ensembles of neurons whose activity was aligned to the mouse’s operant responses (Figures 3D-E). To test whether this was true, we used a k-nearest neighbors (KNN) classifier, which could significantly predict both nose pokes and feeding behavior compared to shuffled data (Figure 3F), while a Gaussian naïve bayes classifier could significantly predict feeding following a left nosepoke (Figure S4D).

Although feeding responses could be reliably decoded by the classifiers,demonstrating that INS^LepR^ ensembles encode operant feeding responses, we found that individual INS^LepR^ cells had heterogeneous responses during the task (Figures 3G-H and S4E), with some neurons being preferentially activated during the nosepoke and some at the start of feeding (Figure 3G), while other cells showed decreased activity once the animal started eating (Figure 3H). We therefore classified the neurons into four distinct clusters with particular response patterns using hierarchical cluster analysis (Figure 3I), and split the data into early, middle and late trials in a session to reflect relative satiation rates from the beginning to the end of the session. In particular, we found one cluster (Cluster 4) that showed high activity preceding feeding events, which was immediately decreased at the start of consumption, similar to Agrp neurons in the arcuate nucleus^43^. However, this cluster grew more pronounced and recruited more cells during the late phase of the task (Figures 3I-J and S4F). This indicates that changes in ongoing satiation level can shift responses towards this specific pattern, which may reflect an overall satiety state of the animal.

INS^LepR^ neurons are specifically responsive to hunger, but not thirst state Because leptin reflects body fat levels and is important for food intake behavior specifically, we next repeated the same calcium imaging experiment, but in a food and water choice task (Figures 4A-C). Mice were food or water restricted and trained to nosepoke for either food or water. After 2-second delay following the nosepoke, they could then freely choose at any point whether they wanted to retrieve food or water.

**Figure 4.**
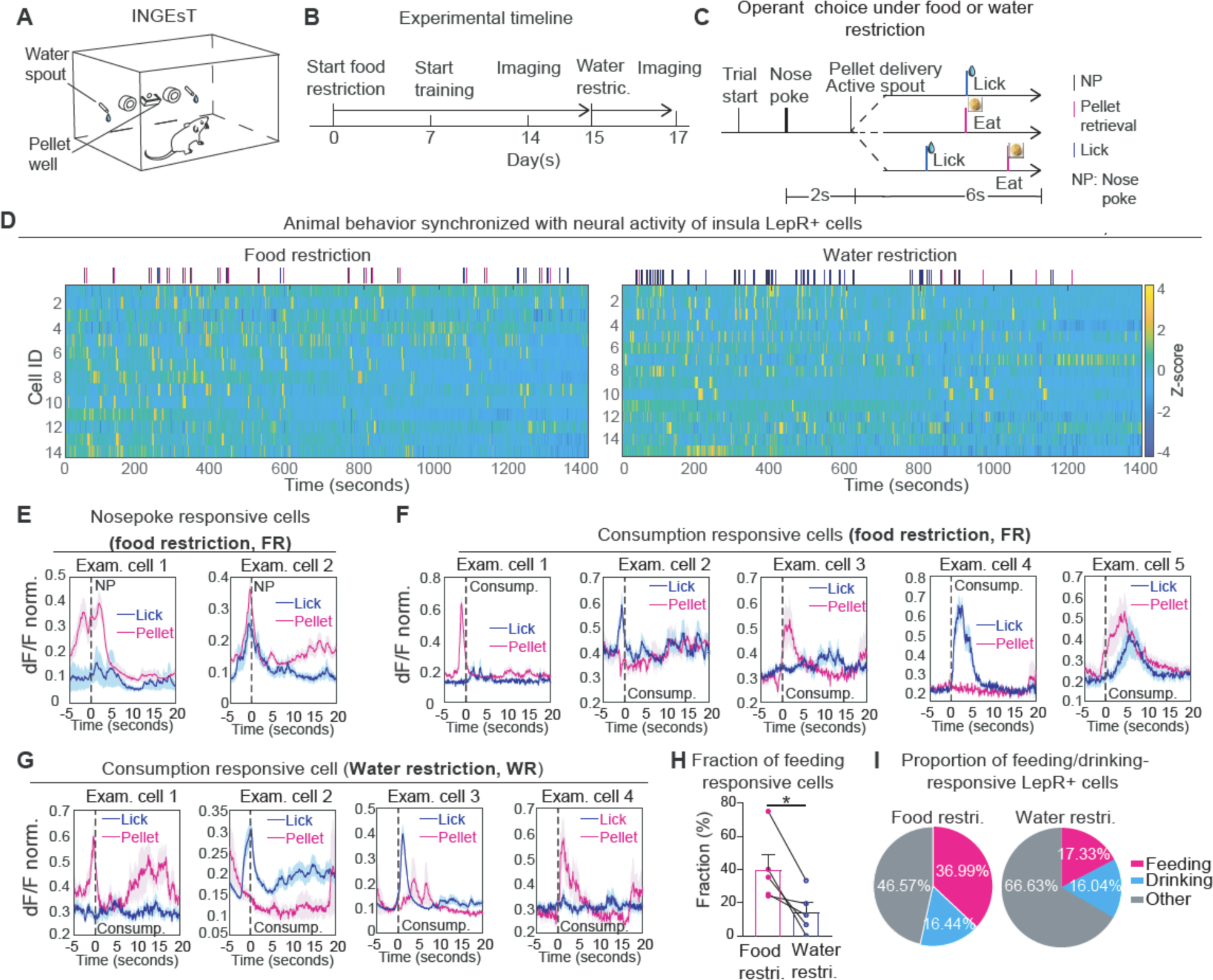
Internal states determine the neural dynamics of LepR+ cells in an operant choice task. (A) Scheme for the INGEsT behavioral chamber. (B) Scheme for operant choice behavioral task. Following the nose poke, a pellet is delivered and the water spout becomes active. Mice may then choose to lick, eat or do both (lick/feed). (C) Example from one mouse of calcium traces (z-score) synchronized to animal behavior in a session under food restriction. Behavior responses are overlayed on top as black (nosepoke), blue (lick) or pink (eat) lines. Each row represents one cell. (D-F) Calcium activity (represented as normalized DeltaF/F (dF/F norm.) in example cells in response to nose poke (**D**) or consumption of food/water (**E-F**) under food (**D-E**) or water (**F**) restriction. Responses to both licking (blue traces) and feeding (pink traces) are shown for each cell. (G) Fraction of feeding responsive cells in individual mice under food or water restriction (n = 5, two-tailed paired t test, * *p* = 0.026). (H) Fraction of feeding/drinking responsive cells pooled from 6 mice under food or water restriction. For more information, see **Figure S5**.

After training, mice were placed on either food or water restriction and then imaged during the choice behavior (Figures 4B-D). For each trial, one of 3 outcomes was available to the mice: 1) nosepoke followed by lick for water (“lick”) 2) nosepoke followed by food pellet retrieval (“feed”) or 3) nosepoke followed by lick and then pellet retrieval (“lick+feed”, Figure 4C). Mice also occasionally retrieved food pellets during an intertrial interval (Figure S5A). In general, mouse behavior was attuned to hunger or thirst state, with feeding prioritized during food restriction sessions and drinking prioritized during water restriction sessions (Figures S5B-E). Interestingly, although lick+feed responses were the least common, they increased non-significantly during food restriction (Figure S5C), suggesting that mice may wet their mouths to prepare for solid food ingestion. It is important to note that despite the preferential response based on internal state, mice did still engage in the opposing response as well, indicating that need states are interdependent. Additionally, this allowed us to investigate responses to feeding and drinking under both food and water restriction. However, the time to lick following a nosepoke was increased during food restriction compared to water restriction (Figure S5E), indicating that the drinking response is qualitatively distinct depending on the need state. We aligned INS^LepR^ neural activity to behavior and examined cell responses under both food and water restriction. Many cells had reliable and predictable responses to specific aspects of the operant behavior, and were tuned to either feeding or licking, but not both (Figures 4E-G, S5F and G). Additionally, comparing cell activity between clusters revealed that INS^LepR^ neural activity was stronger in response to feeding (Figures 4H and I) under food restriction compared to the activity under water restriction. Overall, we found that more cells were feeding responsive under food restriction conditions than water restriction (Figures 4H and I, S4F and G), and we found that INS^LepR^ cells were more highly tuned to feeding-related events compared to licking-related events under food restriction, (36.99% vs. 16.44%). These responses were reduced under water restriction, with less total active cells (Figure 4I), but the responses to drinking remained similar (16.44% to 16.04%), whereas the responses to feeding were decreased 36.99% to 17.33%. Therefore, the activity of INS^LepR^ cells is primarily modulated by feeding behavior, but also by internal state, demonstrating that INS^LepR^ neurons respond to interoceptive information regarding hunger state specifically.

We also carried out calcium imaging during free feeding and drinking, in the absence of an operant response, to assess activity during more naturalistic behavior (Figures 5A and B). In these experiments, mice were given free access to water and food, with a 1-second delay following a lick or pellet retrieval before another reward could be obtained (Figure 5B). In this setup, mice clearly prioritize their immediate need (thirst or hunger) in the first 5 minutes of the session, but then begin to engage in the opposite behavior at the session continues, particularly in the case of food restriction (Figures 5C and D). After aligning the neural activity to the behavior timestamps (Figure 5E-F), responses of individual cells were analyzed (Figures 5G and H, S6A and B). In both cases, cells could be found that were broadly more active during either eating (Figures 5G and H) or drinking (Figure 5H, right), with little to no activity during the other behavior. Examining these individual cells, we found that some cells appeared to be state stable, maintaining their preference for one need state over the other (Figure 5G), whereas some cells were state-switching and shifted their preference depending on the need state (Figure 5H).

**Figure 5:**
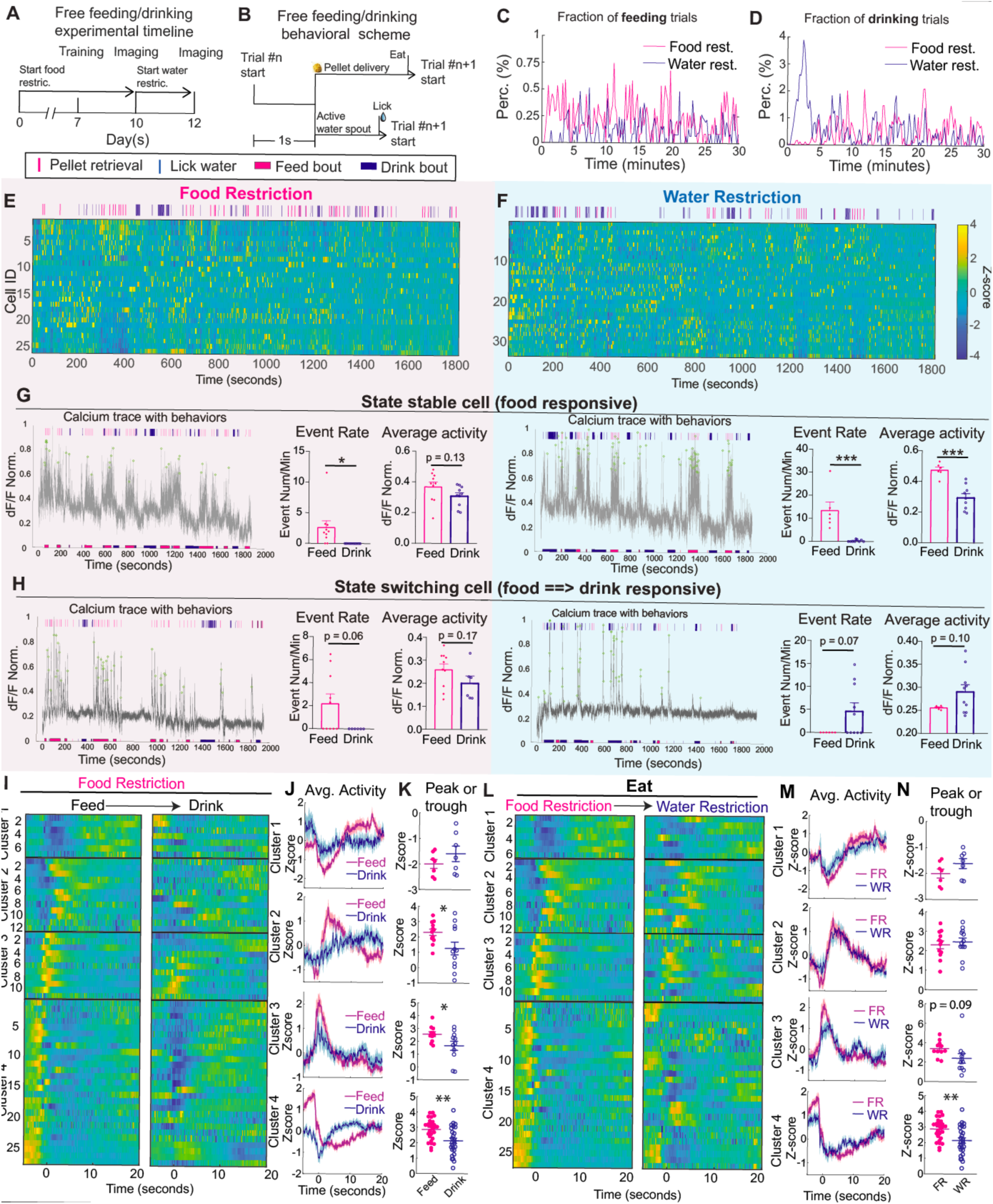
INSLepR neuron activity is food responsive and shaped by internal state. (I) Scheme for feeding/drinking trials. (J) Example from one mouse of calcium traces (z-score) synchronized to animal behavior in one session of the free feeding/drinking task, under food restriction. Behavior is overlaid on top as pink (eat) or blue (lick) lines. Each row represents one cell. (K) Example traces (represented as normalized dF/F) of a feeding responsive cell, which shows stable selectivity under food (**left**) and water restriction (**right**). Individual licks or pellet retrievals are overlaid on top of the traces as pink (eat) or blue (lick) lines. Eating (pink) and drinking (blue) bouts are depicted on the bottom of the trace. Green circles represent detected calcium events. Event rate is represented as the number of events per minute during a food or water consumption bout. The average activity during food or drink consumption bouts is represented as the normalized dF/F of each bout. (n = 10 Feed, n = 10 Drink, two-tailed unpaired t test, * *p* = 0.0223). (Right) Average activity of feeding (magenta) and drinking (blue) bouts (n = 10 Feed, n = 10 Drink, two-tailed unpaired t test, *p* = 0.0129). (L) Example of cell which switches selectivity from food to water under food (**left**) and water restriction (**right**). Individual licks or pellet retrievals are overlaid on top of the traces as pink (eat) or blue (lick) lines. Eating (pink) and drinking (blue) bouts are depicted on the bottom of the trace. Green circles represent detected calcium events. Event rate is represented as the number of events per minute during a food or water consumption bout. The average activity during food or drink consumption bouts is represented as the normalized dF/F of each bout. (n = 10 Feed, n = 6 Drink, two-tailed unpaired t test, * *p* = 0.0634). (Right) Average activity of feeding (magenta) and drinking (blue) bouts (n = 10 Feed, n = 6 Drink, two-tailed unpaired t test, *p* = 0.0168). (M) Classification of insula LepR+ cells in response to animal behavior under food (Left) and water (Right) restriction. (N) Fraction of food-responsive cells which are state stable or state switching between food and water responsive. For more information, see **Figure S6**.

However, examining the specific responses surrounding each need state revealed that, here too, LepR+ cells responded heterogeneously, with peak activity at different times surrounding consumption (Figures 5I and L). We again clustered the neurons using hierarchical clustering and quantified the average activity and the maximum peak or trough to determine whether cells were more broadly modified by need state or behavioral action. We found that activity was strikingly different in most clusters when comparing food vs. water consumption (Figures 5I-K and S6E-G). In contrast, activity was generally similar when comparing consumption during food or water restrictrion, but with slightly larger changes under food restriction (Figures 5L-N and S6H-J). Similarly to the operant based task (Figures 4H and I), there were more total cells modulated during food restriction than under water restriction (34.62% vs 3.85% under food restriction, 21.05% vs. 6.58% under water restriction), but also significantly more cells activated during feeding under food restriction compared to water restriction (34.62% compared to 21.06%, Figure S6C), suggesting that LepR+ cells encode both the internal need state of the animal as well asspecific responses to satisfy hunger. Lastly, we examined the stability of individual responses under food and water restriction. Of the cells that were responsive to feeding under food restriction, 67% remained stably feeding-responsive during water restriction as well, whereas 33% either shifted responses or lost their responsiveness (Figure S6D), again suggesting that INS^LepR^ neurons are responsive to internal state changes.

### Leptin receptors are expressed on both neurons and vascular cells in the insula

A careful examination of our previous in situ hybridization results revealed that there was also LepR+ cells along the pia surface of the cortex (Figure S7A). In order to learn more about this small population of LepR+ cells, we conducted both bulk RNA sequencing from LepR+ polysomes using viral-mediated translating ribosome affinity purification (vTRAP)^44^, as well as single-cell sequencing. We first injected the insula of LepR-Cre mice with a cre-dependent virus expressing EGFP on the ribosomal L10 subunit (Figure 6A). We then pulled down the LepR+ cells with a GFP immunoprecipitation and confirmed successful pulldown via qPCR (Figure 6B). We sequenced the mRNA from both the Input (total insula) and IP (LepR+) populations (Figure 6C, Table S1) and crossed-referenced the top 250 genes with the Allen Institute ABC atlas^45^. We identified that the majority of those enriched transcripts were known to be expressed by vascular cells (Figure 6C, yellow dots). In particular, *Itm2a*, which has been suggested as a potential carrier for blood-brain barrier delivery^46^, marked both the vascular cells as well as neurons in Layer 6 of insula and in the claustrum and had a striking similarity to the LepR expression pattern (Figures S7B-D). In addition, top enriched genes in LepR+ cells of the insula included markers for Layer 6 IT cells and the claustrum, including Nr4a2, Car3 and Gnb4^45^ (Figure 6C).

**Figure 6:**
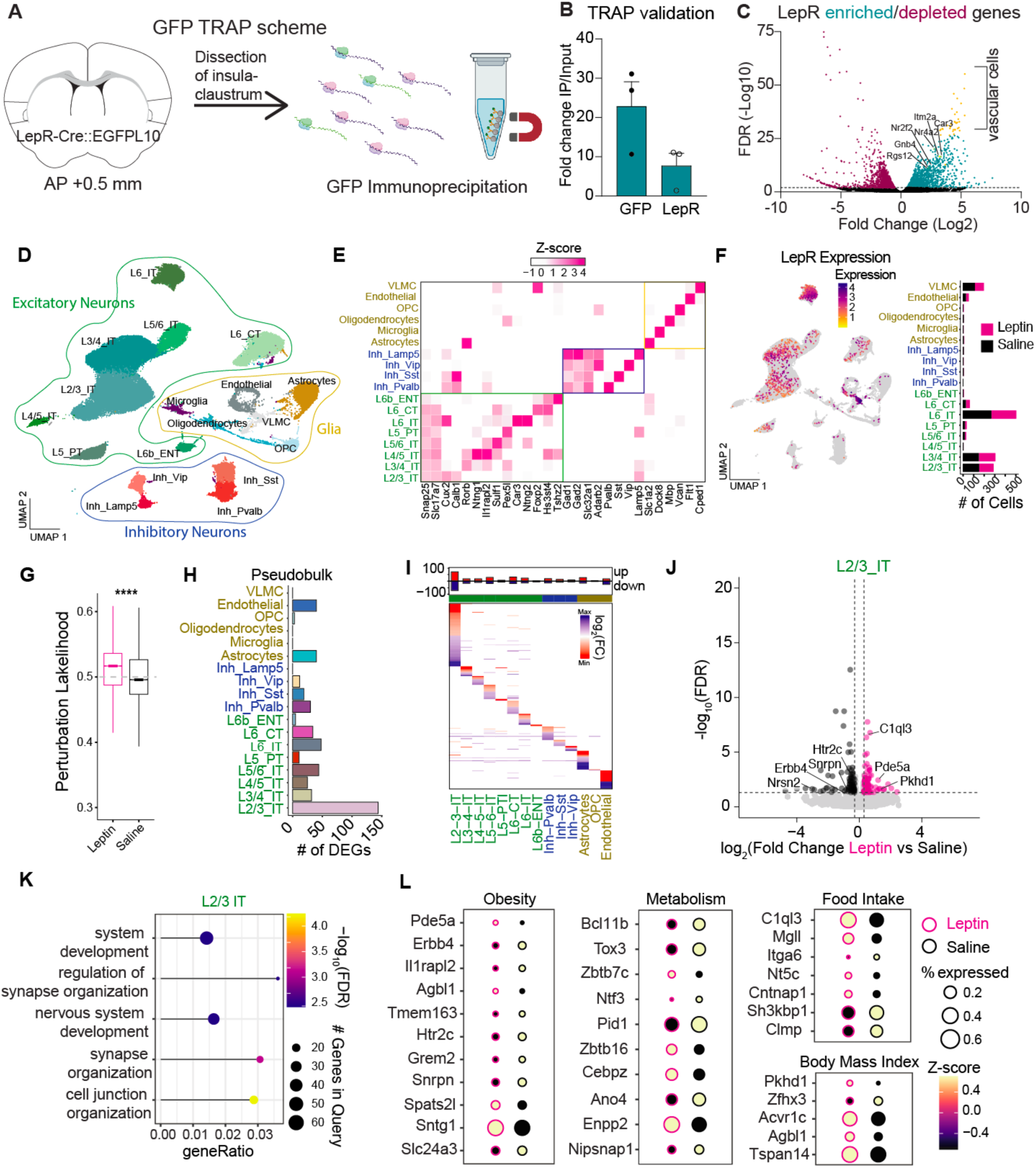
Insular cortex expresses LepR on distinct cell types and transcription profiles are shaped by leptin. (A) Scheme for viral TRAP experiment and GFP immunoprecipitation from the insula of LepR-Cre mice crossed with GFPL10 reporter mouse. (B) Viral TRAP mRNA quality validation of positive controls (Gfp, LepR) run by qPCR. Data is expressed as fold change of the immunoprecipitation fraction (Gfp+/LepR+ cells) compared to input fraction (total insular cortex cells). (C) Plot depicting the false discovery rate (-log10) and average fold change (Log2) of all genes detected during sequencing, comparing the immunoprecipitation fraction to the input fraction. Genes significantly enriched are represented in cyan or yellow. Genes significantly depleted are represented in magenta. Selected genes of interest are highlighted in yellow. Markers of vascular cells are noted by the brackets. (D) Uniform manifold approximation and projection (UMAP) showing plot showing approximately 40,000 nuclei from the insula of 5 saline-treated and 5 leptin-treated mice. Clusters representing excitatory neurons (green), inhibitory neurons (blue) and non-neuronal cells (yellow) are noted. (E) Heatmap showing the normalized expression of representative marker genes, grouped by major cell types. Color indicates the scaled average expression level. Clusters representing excitatory neurons (green), inhibitory neurons (blue) and non-neuronal cells (yellow) are noted. (F) (Left) Expression of LepR+ cells across insular cortex cell clusters. (Right) Number of LepR+ cells in each cluster, divided by saline (control) or leptin-treated mice reveals no difference in LepR+ expression with treatment of leptin. (G) Significant change in perturbation likelihood by leptin treatment across insular cortex cells (Left). Perturbation likelihood is also depicted by individual clusters (Right). Significance determined by two-sided Wilcoxon Rank Sum test (p < 0.0001) (H) Barplot of differentially expressed genes (DEGs) in the distinct cell-type following leptin treatment (Pseudobulk, DESeq2; FDR < 0.05, abs(log2(Fold Change)) > 0.3). (I) Heatmap depicting the DEGs in the distinct cell-types ranked by fold change difference between leptin treated mice and control mice (blue: enriched, red: depleted). (J) Volcano plot depicting significantly enriched (pink) and depleted (grey) genes in insular cortex Layer 2/3 cells following leptin treatment. (K) Bubble chart highlighting the top 5 functional categories enriched in DEGs of L2/3 cells. Gradient color corresponds to the −log10(FDR), dot size correspond to the number of DEGs in the functional category. (L) Expression level (z-score) and percentage of cells expressing differentially expressed genes in layer 2/3 following leptin treatment. Genes are grouped into previously known functions related to food intake, obesity, metabolism and body mass index. For more information, see **Figures S7 and S8**.

### Leptin leads to distinct gene expression changes in the insula

We next examined whether peripheral leptin infusion would lead to downstream changes in insula gene expression using single nuclei sequencing. Mice were injected with either saline of leptin (5 μg/g of body weight) and were euthanized 24 hours later, followed by nuclei isolation and sequencing. Insula nuclei could be clustered into 17 different clusters (Figure 6D), defined by specific markers which indicated their overall cell identity (Figure 6E). We first examined the expression of leptin receptors within the insula itself, which confirmed our bulk sequencing results that LepR are present both in layer 6 IT neurons and in vascular cells, particularly the VLMC and endothelial populations (Figure 6F). Previous reports have demonstrated that leptin can be actively transported across the blood brain barrier, and knockout of LepR in brain endothelial cells leads to changes in feeding and body weight^47–49^. Together, these results suggest that leptin receptors are present on vascular cells in the insula and likely transport leptin through the blood brain barrier directly into the cortex.

We then used a MELD^50^ to determine how leptin impacted the cell types in the insular cortex. Notably, we observed a significant overall and cell class specific perturbation caused by leptin (Figures 6G and S8A-C). Interestingly, the leptin response revealed a robust effect in multiple neuronal cell types, such as L2/3 IT and L3/4 IT, as well as in cell types related to the blood-brain barrier, including VLMCs and endothelial cells (Figures 6G and S8D). These results were confirmed by AUGUR^51^, which further validated the leptin-induced changes across these cell types (Figure S8E-F). To gain a deeper understanding of how leptin induces transcriptional changes acrossinsular cortex cell types, we employed a pseudobulk differential expression approach, which offers increased robustness by aggregating signal across cells within each type. We identified hundreds of differentially expressed genes (DGE), with more than 150 genes belonging to layer 2/3 IT neurons (Figures 6H-J), which are a population of neurons that have previously been shown to be involved in regulating food intake, particularly in associative learning contexts^19,20^. To investigate the biological function of leptin-induced differentially expressed genes (DEGs) in L2/3 IT neurons, we performed gene set enrichment analysis using scToppR^52^. This analysis revealed that leptin-induced transcriptional changes are enriched for genes involved in synaptic function and neuronal development (Figure 6K). We next asked whether leptin also modulates the expression of genes linked to feeding behaviors. Interestingly, among these genes (Figure 6L, Table S2) were approximately 20% that have been associated with obesity and diabetes (*Pde5a, Erbb4, Il1rapl2, Agbl1, Tmem163, Htr2c, Grem2, Snrpn, Spats2l, Sntg1, Slc24a3*), human body mass index (*Pkhd1, Zfhx3, Acvr1c, Tspan14*), mouse food intake and body weight (*C1ql3, Mgll, Itga6, Nt5c, Catnap1, Sh3kbp1, Clmp*) and metabolism (*Bcl11b, Tox3, Zbtb7c, Ntf3, Pid1, Zbtb16, Cebpz, Ano4, Enpp2, Nipsnap1*), both in the brain and in the periphery. Taken together, these findings suggest that leptin orchestrates a network of transcriptional changes within the insular cortex, primarily targeting a population of neurons involved in regulating food intake, thereby contributing to the modulation of feeding behavior.

### Leptin reshapes insular cortex population activity to feeding through local connections

Our bulk and single-cell sequencing data (Figure 6) showed that neuronal LepR+ cells are expressed mainly in Layer 6 IT Car3+ neurons. A recent study showed that Car3 neurons are both transcriptionally and morphologically distinct from other cortical glutamatergic cells, projecting mainly within the cortex or to ipsilateral cortex^45,53^.

Morphologically these neurons have widely dispersed fibers compared to Layer 6 CT cells which have a characteristic apical dendrite. Similarly, we traced the morphology of both LepR+ and LepR-cells in the insula and found that INS^LepR^ have widely dispersed fibers compared to LepR-neurons examined (Figure 7A), with a greater number of intersections close to the soma (Figure 7B). Because layer 6 IT cells are known to project primarily intracortically, we examined local connections (Figures S9A-E) and found that INS^LepR^ were connected to ∼20% of patched cells (Figure 7C-D). We therefore hypothesized that leptin would reshape local insular cortex activity to modulate food intake. Wild-type C57BL/6J mice were injected with synapsin-driven GCaMP8m and a GRIN lens implanted to image from general insula neuronal populations (Figure 7E). We then repeated the free-feeding and drinking experiment (Figure 5), but with injections of leptin or PBS given 12 hours before the imaging session (Figures S9G-L). This revealed a similar imaging pattern, as well as neuronal clusters as we previously observed (Figures 5G-I and S6H-J). Although some clusters showed a similar response between PBS and Leptin treatment, population activity with PBS treatment revealed stronger changes than the activity under leptin treatment (e.g. Figure S9I, Cluster 3; Figure S9L, Clusters 3 and 4). We then reduced dimensions of the INS^LepR^ population activity in response to food or water consumption, with and without leptin (Figure 7F) and projected the neural activity around pellet retrievals and licks in principal component subspace. In all four cases, PC1 accounted for over 86% of the total variance, and first three PCs together accounted for over 96% of the variance. We found that food and water consumption had distinct neural activity trajectories under normal conditions (Figures 7F-H), which we quantified using the dissimilarity analysis, procustes distance and angle between subspaces, where the higher value means lower similarity.

**Figure 7.**
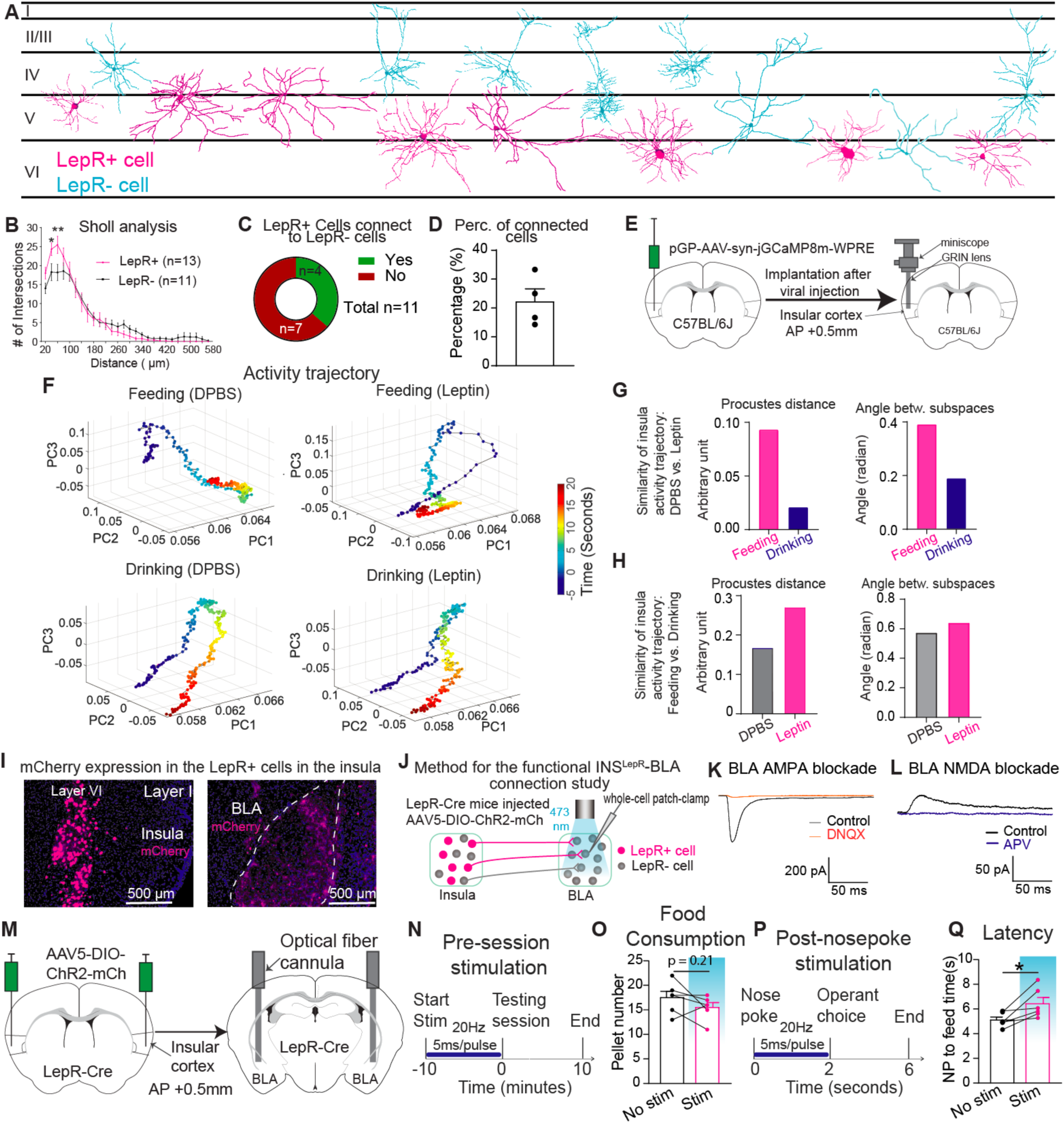
Stimulation of leptin receptors shapes neural dynamics in the insula. (A) Morphology of LepR+ (magenta) and LepR- (cyan) cells in the insula, across cortical layers. (B) Sholl analysis for the morphological comparison between LepR+ (magenta) and LepR- (cyan) cells. Data is represented as the number of intersections at different distances from the soma, measured in μm. (n = 13 LepR+, n = 11 LepR-, two-way ANOVA followed by Turkey post-hoc test, F_(1,616)_ = 0.0108; at 40 μm, * *p* = 0.0268; at 60 μm, ** *p* = 0.002). (C) Percentage of connected cells around the patched cells. 22.3% of cells around the patched cells (4 cells from **E**) can be activated. (D) Scheme for the method of viral injection and GRIN lens implantation to image calcium activity in the general insular cortex population following saline or leptin treatment. (E) Trajectory of activity during feeding or drinking bouts 12 hours after PBS (control) or leptin (5 μg/kg) administration using principal component analysis (PCA) . Data is shown as activity over time from 5 seconds before to 20 seconds after the bout begins. (F) Similarity of insula activity trajectories at feeding/drinking between DPBS- and leptin-treated group. Top: Procustes analysis (feeding, magenta; drinking, blue). Bottom: Subspace angle analysis (feeding, magenta; drinking, blue). (G) Similarity of insula activity trajectories in DPBS- and leptin-treated group between feeding and drinking behavior. Top: Procustes analysis (leptin, magenta; DPBS, blue). Bottom: Subspace angle analysis (leptin, magenta; DPBS, blue). (H) Representative images of terminal projection mapping from mCherry expression in LepR+ cells in the insular cortex injection site (left) and in axon terminals in the basolateral amygdala (right). (I) Scheme for the method of studying functional connectivity by using whole-cell patch-clamp recording. (J) Excitatory postsynaptic current (EPSC) is inducted by 2 ms of blue light (Opto-EPSC). (K) Scheme for viral injection and optical fiber cannula implantation for optogenetic stimulation of BLA-projecting LepR+ cells in the insula. (P) Scheme for pre-session optogentic stimulation. (Q) Pellet number consumed over one testing session with pre-session stimulation (magenta) or control (blue)) (n = 6 Stim, n = 6 No stim, two-tailed paired t test, *p* = 0.2098). (R) Scheme for the protocol of post-nosepoke stimulation. (S) Latency to pellet retrieval following nose poke with stimulation (magenta) or control (blue) (n = 6 Stim, n = 6 No stim, two-tailed paired t test, * *p* = 0.0276). For more information, see **Figures S9-S10**.

Interestingly, following leptin administration, the trajectory to food consumption was dramatically shifted compared to drinking (Figure 7G), indicating that leptin shapes population activity to food consumption more than water consumption, in the insula. In addition, leptin increased the dissimilarity of the trajectories between feeding and drinking when compared to the PBS control. Overall, leptin treatment shaped the insula dynamics in response to feeding and drinking behavior.

### INS^LepR^ neurons project to BLA and decreases learned feeding

In contrast to the overall population of Car3 neurons, which make projections to ipsilateral cortex, LepR+ neurons appear to make connections primarily within the ipsilateral insula itself. We found only one projection to the basolateral amygdala, using both terminal tracing (Figures 7I and S10A) and a color-switching retrograde virus injected into the BLA (Figure S10B and C). We conducted patch clamp electrophysiology to confirm connections between INS^LepR^ cells and both insula and basolateral amygdala neurons (Figure 7J-L, and S10C-I), finding that these connections are glutamatergic and that there is a lower probability of neurotransmitter release and lower NMDA/AMPA ratio at the BLA compared to local connections. Together, these results suggest that INS^LepR^ neurons act primarily to integrate leptin signal and send internal state information primarily locally, but also to the basolateral amygdala.

We therefore tested whether activating INS^LepR^ terminals at the basolateral amygdala could replicate the optogenetic effects with cell body activation (Figure 7M). We found that in contrast to our previous results, activation of BLA terminals did not significantly change the pellet retrieved following a nosepoke (Figures 7N and O), but did increase the time between nosepoke to pellet retrieval (Figures 7P and Q). This indicates that activation of BLA terminals has a specific effect on the timing to retrieve a food reward, whereas the insula connections may impact the likelihood of retrieving pellets.

Taken together, our results point to a strong role of leptin and leptin receptors in the insula in providing direct interoceptive information that guides learned feeding behaviors, through local connections within the insula and a projection to the basolateral amygdala.

## DISCUSSION

Interoception, re-defined by Bud Craig more than 2 decades ago as a bridge between the physical and emotional and an awareness of the bodily self^10,12^, is now considered to be a primary function of the brain in coordinating bodily responses and making decisions. In Craig’s conception, the insula is a key site for interoception and integrating internal states with exteroceptive information. However, nearly all the evidence to support this hypothesis has come from human brain imaging studies and from the anatomical perspective of polysynaptic connections between the vagus nerve and the insula. In contrast, interoceptive signals directly from the body, such as leptin, ghrelin and glucagon-like peptide-1, are thought to act at sites that do not have an intact blood brain barrier, such as the arcuate nucleus and other areas of the hypothalamus and hindbrain. Here, we make a number of significant advances from the established dogma: 1) We show that the insula receives direct interoceptive input in the form of leptin, which acts on leptin receptors in the insula. 2) We find that LepR is present not just on neurons, but also on vascular cells, in the insula, pointing to a possible active transport through the BBB as a mechanism for entry into the cortex. 3) We find that interoceptive signals delivered to the insula serve primarily to deliver information about bodily state that can be used to shape neural activity to food that is used for reward-based learning rather than homeostatic control of feeding. This work, therefore, makes fundamental advances in understanding the function of the insula and the neural basis of interoception, which opens up new lines of investigation across a number of different fields.

Through a combination of morphology tracing, single-nuclei and bulk RNA sequencing, we found that INS^LepR^ neurons are expressed on Car3 positive neurons of Layer 6 and the claustrum. This population is a recently described class of neurons in the isocortex with a unique morphological and transcriptomic pattern^45,53^. Interestingly, cortical Car3 neurons are transcriptionally similar to claustrum neurons, but have a much more restricted intracortical projection pattern. Based on our results, INS^LepR^ appear to be more similar to cortical Car3 neurons, as they project mainly within the insula itself and not widely throughout the cortex. However, both cortical and claustrum Car3 neurons were shown not to have collateral projections to the striatum, whereas INS^LepR^ neurons project to the BLA, suggesting that INS^LepR^ may be a distinct subpopulation of this class of neurons in the insula, rather than just a general Car3+ marker. Another recent study described an atlas of the claustrum and insula region, demonstrating that the boundaries between the insula and claustrum are not clear, and proposing to distinguish between cell types rather than strict anatomical boundaries^54^. Thus, INS^LepR^ may represent a single population of neurons distributed in this more ambiguous insula-claustrum area. Our finding that leptin receptors are present not just on neurons, but also on vascular cells is also supported by previous literature. Previous whole-brain single-cell sequencing identified LepR on VLMC cells^45^, and deletion of LepR in brain endothelial cells prevented leptin uptake in the cortex as well as ventral tegmental area, and led to hyperphagia and weight gain on high fat diet, but not normal chow^49^. This aligns with our results indicating that, in the insula, LepR does not mediate homeostatic feeding, and suggests that blood-brain barrier transport of LepR may be necessary to impart interoceptive signals used for non-homeostatic feeding, but not for baseline energy balance demands.

Murayama et al^38^ reported that leptin enhances inhibitory synaptic transmission from Vgat+ neurons to pyramidal excitatory neurons in the rat insular cortex and proposed that LepR might be expressed on insula Vgat+ neurons. However, both our *in situ* hybridization and single nuclei sequencing data confirmed INS^LepR^ are exclusively present in the glutamatergic neurons, and not in the inhibitory neurons in mice. This was also confirmed by examination of the Allen Institute ABC Atlas^45^. We found that leptin increases intrinsic excitability in glutamatergic INS^LepR^ cells, but not in LepR-cells in the insula. Thus, it is possible that the enhancement of uIPSCs from insula Vgat+ cells to pyramidal cells observed by Murayama et al. might be due the leptin’s effects on glutamatergic INS^LepR^ cells connecting to LepR-Vgat+ cells. It has long been appreciated that feeding is a complex motivated behavior and that feeding responses are coordinated by a diverse set of brain regions and under a variety of conditions.

However, the underlying assumption has been that homeostatic brain regions control the simple homeostatic aspect of food intake through hormonal signaling from the body, whereas the more complex aspects of food intake are controlled by higher-order brains regions in the limbic and cortical areas. Our study indicates, instead, that hormonal signals can also influence these higher order brain areas, and that the insula in particular receives interoceptive signals and integrates that information within the insula. Notably, we find that INS^LepR^ neurons do not form many connections outside of the insula, with only a single projection to the basolateral amygdala detected. Thus, it appears that these cells act primarily locally to receive internal state information and relay it to other insula neurons. Accordingly, we find that leptin administration primarily impacts neurons in Layer 2/3 of the cortex where the most DEG’s were found. This is consistent with previous literature indicating that insula layer 2/3 neurons play a critical role in food-associative learning^19,20^. These results underscore the importance of integrating interoceptive input within the insula in order to organize top-down control of food intake. Moreover, the effects of leptin administration and INS^LepR^ activation occur over relatively long time scales. A single injection of leptin in the insula significantly modifies food intake and body weight for two days, and activation of INS^LepR^ before testing in an operant task leads to a sustained decrease in pellet retrieval over the subsequent 10 minute session. These timescales are consistent with the idea that leptin is a long-term signal of total body fat levels, rather than a short term, meal termination signal, and suggests that leptin levels are utilized in the insula over longer timescales as well.

We found the INS^LepR^ neural responses to feeding and drinking are shaped by internal state and that INS^LepR^ neural dynamics are modulated by changes in internal state. Previous studies compared the neural dynamics in the insula between a food restricted and sated state^19,25^. Because mice naturally do not exhibit eating behavior while sated, it is not possible to observe different neural responses to feeding under different conditions. To overcome this, we developed a free feeding and drinking paradigm and an operant food and water choice task to investigate neural dynamics under both hunger and thirst states^42^. Notably, mice still engage in feeding bouts under water restriction, albeit with less frequency and later in the session, so that we can examine responses to feeding under both conditions. We observed more feeding responsive INS^LepR^ cells under hunger states than thirst states, and the amplitude of feeding response was higher under hunger state compared to thirst state. In addition, our data showed that leptin’s effect on feeding was dependent on internal state. Local infusion of leptin into the insular cortex produced an inhibition of feeding for 48 hours in mice without food restriction, but the same treatment produced an inhibition of feeding for around 24 hours in mice under food restriction. Murayama et al^38^ also reported that rat insula excitability shaped by leptin infusion was dependent on the internal energy states. Thus, our data provides substantial evidence that internal states shape the INS^LepR^ neural response and associated behaviors.

We observed that chemogenetic manipulation or ablation of LepR had no effect on homeostatic food intake or body weight. These results are consistent with those reported in the lateral hypothalamus (LH), in which the ablation of LepR resulted in no body weight change^55^, although studies using optogenetic activation in the LH have had inconsistent results^56–60^. In addition, the authors observed a decrease in appetitive reward learning in the LH, consistent with our results that optogenetic activation decreased operant-based pellet retrieval. In addition, our calcium imaging results are also consistent with those in the lateral hypothalamus, in which LepR+ neurons have heterogeneous responses throughout the test session to distinct aspects of the task.

Another study demonstrated that LH LepR neurons can be subdivided into food-seeking and food-consuming subpopulations^57^. Although our task was not designed to specifically test for this, our results are also consistent with such a distinction.

Previous studies have examined the role of insular cortex projections to the BLA. One study found that activation of the anterior insula to BLA pathway after novel taste consumption induced subsequent aversion to that tastant^61^, whereas activation of posterior insula to BLA decreased water intake^62^. In another study, both activation and inhibition of anterior insula to BLA projecting neurons were shown to decrease anxiety-like behavior^63^. These studies underscore the utility of studying a molecularly defined population like INS^LepR^ neurons, which make up only a portion of the BLA projection from the insula, as it is possible that conflicting results arise from studying a mix of projecting cell types.

The current study demonstrates a role of LepR in direct interoceptive sensing by the insula. This suggests that there are likely other populations which mediate other aspects of interoception. A recent study reported that Glp1r+ neurons in the insula regulate feeding^64^, however Glp1 is released both from peripheral and central sites. It is therefore not yet known whether insula Glp1r detects peripheral internal state signals.

Another recent study demonstrated that insula neurons track thirst state independent of subfornical organ stimulation^25^, which promotes drinking. Thus, it is possible that the insula also has direct interoceptive input into thirst states, which have yet to be uncovered. Moreover, interoceptive states reflecting positive and negative valence emotions are likely also encoded in the insula, and whether this aspect of interoception would also be mediated by molecularly defined subsets of cells is not known.

### Limitations of the study

In the current study we describe the role of INS^LepR^ in regulating short-term food intake using acute optogenetic activation, but injections of leptin directly into the insula suggest the capacity to promote global changes in food intake or body weight. Future investigations would be needed to understand if manipulations of leptin receptors would have more long-term effects relating to obesity or other food-intake disorders.

In addition, our transcriptomic analysis concluded that leptin significantly altered gene expression in Layer 2/3 neurons of the insula, but we did not investigate the role of these gene and their associated cells in regulating food intake. A comprehensive understand of how food intake is regulated by the insula will require a knowledge of how these gene expression changes relate to changes in feeding.

Lastly, although our results corroborate previous findings of blood-brain barrier transport of leptin into the brain, the exact mechanisms by which this occurs in different brain regions are still unclear. In the hypothalamus, tanycytes appear to regulate active transport^65^, while in the ventral tegmental area, brain endothelial cells were shown to mediate leptin entry^49^. In our study we demonstrate LepR presence on yet another population, VLMC cells. A more complete understanding of this process will be beneficial for our understanding of leptin action in the brain, as well as for clinical applications.

## Supporting information

S1 Video ObRb Insula

SI Video 2 Activity and Behavior

## RESOURCE AVAILABILITY

### Lead contact

Requests for further information and resources should be directed to and will be fulfilled by the lead contact, Sarah Stern.

### Materials availability

This study did not generate new, unique reagents

### Data and code availability

Sequencing data will be deposited to GEO database before publication. Code will be made available on Github prior to publication.

Accession numbers and links will be available and placed here prior to publication. •Any additional information required to reanalyze the data reported in this paper is available from the lead contact upon request.

## ACKNOWLEDGMENTS

The authors thank Colleen Neiner, Jose Salazar Gamboa, Amanda Coldwell, Dr. Idris El-Amin and the staff of the MPFI Animal Resource Center. We thank Nicole Daniel and Abraham Rivera from the MPFI Mechanical Workshop for custom behavioral chambers. We thank Nicolai Urban, Naomi Kawasawa and Jenny Wu for assistance with light microscopy and molecular assays. We thank Robert M. Witwicki and Li Pan of the UF Scripps Genomics Core, as well as Gogce Crynen of the UF Scripps Bioinformatics Core, for their contributions to RNA sequencing data and analysis, respectively. We thank Timothy Holford for discussing and providing advice on patch-clamp recording experiments. We thank Jeffrey Friedman for assistance with LepR-Cre:TdTomato mice. We thank Lin Tian, David Fitzpatrick, Estefania Azevedo, Federica Cruciani and Sebastien Bullich for helpful discussions and comments on the manuscript. Some schemes were made using Biorender. This work was supported by an NIH R00 (SAS), Brain Research Foundation Seed Grant (SAS), One Mind Foundation (SAS), an NIH New Innovator DP2 (SAS) and the Max Planck Society (SAS). S.S., B.G., and S.B. are supported by the CNDD Genomics and Bioinformatics Core (P20 GM148302).

## AUTHOR CONTRIBUTIONS

Conceptualization: Z.Z and S.A.S.; Behavior: Z.Z., S.A., E.M.; Viral-TRAP: Z.Z. and S.A.S. Single-cell experiment: S.A.; Analysis/bioinformatics: Z.Z., B.X., S.S., B.G., and S.B.; Electrophysiology: Z.Z.,C.V.W., and M.M.B.; Imaging and histology: Z.Z., I.M., A.S., Z.L., and E.M; Morphology tracing: Z.Z., and M.K.; writing-original draft: Z.Z. and S.A.S; Funding acquisition: S.A.S; Supervision: M.M.B, S.B. and S.A.S.

## DECLARATION OF GENERATIVE AI AND AI-ASSISTED TECHNOLOGIES

Generative AI and AI-assisted technologies were not used.

## SUPPLEMENTAL INFORMATION

**Document S1:** Figures S1-S10

**Table S1: LepR Viral TRAP**

**Table S2: DEG analysis – single-nuclei sequencing**

**Video S1: LepR mCherry in insula**

**Video S2: LepR calcium activity**

## STAR ★ METHODS

### KEY RESOURCES TABLE

| REAGENT or RESOURCE | SOURCE | IDENTIFIER |
| --- | --- | --- |
| **Antibodies** |  |  |
| Mouse monoclonal anti-GFP (Heintz Lab TRAP anti-GFP 19F7) | MSKCC Antibody & Bioresource Core Facility | Cat#Htz-GFP-19F7; RRID: AB_2716736 |
| Mouse monoclonal anti-GFP (Heintz Lab TRAP anti-GFP 19C8) | MSKCC Antibody & Bioresource Core Facility | Cat#Htz-GFP-19C8; RRID: AB_2716736 |
| Anti-mCherry antibody | abcam | Cat#ab205402; RRID: AB_2722769 |
| Goat anti-Chicken secondary antibody | Thermo Fisher Scientific | Cat#A-11042; RRID: AB_2534099 |
| **Bacterial and virus strains** |  |  |
| pAAV-hSyn-DIO-mCherry | Addgene | Addgene AAV5; 50459-AAV5 |
| pAAV-hSyn-DIO-hM3D(Gq)-mCherry | Addgene | Addgene AAV5; 44361-AAV5 |
| AAV-hSyn-DIO-hM4D(Gi)-mCherry | Addgene | Addgene AAV5; 44362-AAV5 |
| pAAV-EF1a-double floxed-hChR2(H134R)-mCherry-WPRE-HGHpA | Addgene | Addgene AAV5; 20297-AAV5 |
| pOTTC1032 - pAAV EF1a Nuc-flox(mCherry)-EGFP | Addgene | Addgene AAVrg; 112677-AAVrg |
| pGP-AAV-syn-FLEX-jGCaMP8m-WPRE | Addgene | Addgene AAV1; 162378-AAV1 |
| pGP-AAV-syn-jGCaMP8m-WPRE | Addgene | Addgene AAV1; 162375-AAV1 |
| **Biological samples** |  |  |
| Chemicals, peptides, and recombinant proteins |  |  |
| Recombinant Mouse Leptin Protein | R&D Systems | 498-OB-01M |
| Clozapine-N-oxide | Tocris | Cat#4936; CAS: 34233-69-7 |
| Dulbecco's phosphate-buffered saline (DPBS) | Gibco | Cat# 14190136 |
| 1,2-diheptanoylsn-glycero-3-phosphocholine (DHPC) | Avanti Polar Lipids | Cat#850306; CAS: 39036-04-9 |
| Tetrodotoxin Citrate (TTX) | Tocris | Cat#1069; CAS: 18660-81-6 |
| DL-AP5 Sodium Salt | Tocris | Cat#3683; CAS: 338404-52-7 |
| NBQX disodium Salt | Tocris | Cat#1044; CAS: 479347-86-9 |
| (-)-Bicuculline methobromide | Tocris | Cat#0131/10; CAS: 38641-83-7 |
| 4-Aminopyridine (4-AP) | Tocris | Cat#0940; CAS: 504-24-5 |
| Picrotoxin | Sigma-Aldrich | Cat#P1675; CAS: 124-87-8 |
| Choline Chloride | Sigma-Aldrich | Cat#C7017-100g; CAS:67-48-1 |
| Sodium Pyruvate | Sigma-Aldrich | Cat#P2256-100g; CAS:113-24-6 |
| Sodium Chloride | Sigma-Aldrich | Cat#S5886-1Kg; CAS:7647-14-5 |
| Thiourea | Sigma-Aldrich | Cat#T7875-100g; CAS:10450-60-9 |
| Sodium Ascorbate | Sigma-Aldrich | Cat#A7631-100; CAS:134-03-2 |
| Sodium Bicarbonate | Sigma-Aldrich | Cat#S5761-1kg; CAS:144-55-8 |
| Sodium Phosphate monobasic | Sigma-Aldrich | Cat#S5011-500g; CAS: 7558-80-7 |
| Calcium Chloride dihydrate | Sigma-Aldrich | Cat#C7902-500g; CAS:10035-04-8 |
| Magnesium Chloride hexahydrate | Sigma-Aldrich | Cat#M2393-500; CAS:7791-18-6 |
| Potassium Chloride | Sigma-Aldrich | Cat#P5405-500g; CAS:7447-40-7 |
| Glucose | Sigma-Aldrich | Cat#G7021-1kg; CAS:50-99-7 |
| HEPES | Sigma-Aldrich | Cat#H4034-100g; CAS:7365-45-9 |
| EGTA | Sigma-Aldrich | Cat#E4378-100g; CAS:67-42-5 |
| Potassium Gluconate | Sigma-Aldrich | Cat#P1847-500g; CAS:299-27-4 |
| CsOH 50% solution | Sigma-Aldrich | Cat#232041-50g; CAS:21351-79-1 |
| Gluconic acid 50% solution | Sigma-Aldrich | Cat#G1951-25g; CAS:526-95-4 |
| Adenosine 5'triphosphate Magnesium Salt | Sigma-Aldrich | Cat#A9187-1g; CAS: 74804-12-9 |
| Guanosine 5'triphosphosphate | Sigma-Aldrich | Cat#G8877-250mg; CAS:36051-31-7 |
| Biocytin | Sigma-Aldrich | Cat#B4261; CAS: 576-19-2 |
| Pierce™ Recombinant Protein L, Biotinylated | Thermo Fisher Scientific | Cat#29997 |
| HEPES, 1M Solution, pH 7.3 | Thermo Fisher Scientific | Cat#J16924.K2; CAS:7365-45-9 |
| DL-Dithiothreitol (DTT) | Sigma-Aldrich | D9779-5G; CAS: 3483-12-3 |
| Cycloheximide | Sigma-Aldrich | C7698-5G; CAS: 66-81-9 |
| Spermine | Sigma-Aldrich | S11471-1G; CAS: 306-67-2 |
| Spermidine | Sigma-Aldrich | S2501-1G; CAS: 334-50-9 |
| Critical commercial assays |  |  |
| Deposited data |  |  |
| Viral-TRAP Data | The Max Planck Florida Institute | TBD |
| ScRNA-Seq Data | The Max Planck Florida Institute | TBD |
| Experimental models: Organisms/strains |  |  |
| Mouse: C57BL/6J | The Jackson Laboratory | RRID:IMSR_JAX:000664 |
| Mouse: ObRb-Cre | The Jackson Laboratory | RRID:IMSR_JAX:008320 |
| Mouse: EGFP-L10a | The Jackson Laboratory | RRID:IMSR_JAX:022367 |
| Software and algorithms |  |  |
| GraphPad Prism 8.0 | GraphPad Software | N/A |
| ImageJ | NIH | N/A |
| Imaris | Imaris, Oxford instrument, UK | N/A |
| Matlab R2022a | MathWorks | N/A |
| Python 3.11 | Python | N/A |
| Other |  |  |
| Confocal microscope | Zeiss | LSM780 |
| Epifluorescence microscope | Keyence | BZ X800 |
| Stereotaxic apparatus | KOPF instruments | Model 900 |
| Micropump | Harvard Apparatus | Pump 11 Picro Plus Elite |
| Guide-Cannula (Length 4.8 mm) | Plastic One | C315GMN/SPC |
| Internal-Cannula (Length 5.1 mm) | Plastic One | C315LIMN/SPC |
| Osmotic pump | ALZET | 2002 |
| Brain infusion kit | ALZET | Cat#0008663 |
| Electrophysiology Rig | Zeiss | Zeiss AXIO Examiner.D1 |
| Fiberoptic cannula | Doric Lenses | MFC_200/245-0.37_4mm_ZF1.25_FLT |
| Bioanalyzer | Agilent | Cat#2100 |
| Vibratome | Leica | VT1000S |
| Cryostat | Leica | CM3050 S |
| Superfrost Plus Slides | Fisher Scientific | Cat#22-037-246 |
| RNAscope® Multiplex Fluorescent Detection Kit v2 | ACD | Cat# 323110 |
| HybEZ Humidity Control Tray | ACD | Cat#PN 310012 |
| EZ-Batch Wash Tray | ACD | Cat# PN 321717 |
| HybEZ Oven | ACD | Cat# PN 321710 |
| Leptin receptor probe | ACD | Cat#402731-C2 |
| VGAT probe | ACD | Cat#319191-C3 |
| VGLUT1 probe | ACD | Cat#416631-C3 |
| High-throughput sequencing platform | Illumina | Nextseq-2000 |
| Nalgene™ PPCO Conical-Bottom Centrifuge Tubes | Thermo Fisher Scientific | Cat#3103-0050PK |
| Stopper rubber | Eisco | Cat#CH0314K1H |
| Drinking tube (2.5") | Ancare | Car#TD-100 |
| Miniscope | Inscopix | nVue |
| ProView™ Integrated Lens 0.5mm x 6.1mm | Inscopix | Cat#1050-004415 |
| 470 LED | THORLabs | M470F4 |
| Fluoromount-G™ Mounting Medium, with DAPI | Thermo Fisher Scientific | Cat#00-4959-52 |
| Trypan blue | Gibco | Cat#15-250-061 |
| Hemocytometer | Bright-Line | Cat#Z359629-1EA |
| Brightfield microscope | OLYMPUS | CKX41 |
| RNeasy Mini Kit | QIAGEN | Cat#74104 |
| RNase Zap | Thermo Fisher Scientific | Cat#AM9782 |
| Complete Mini, EDTA-free Protease Inhibitor | Roche | 4693124001 |
| GEM blade | Electron Microscopy Sciences | Cat#71970-WA |
| UltraPure™ BSA (50mg/mL) | Invitrogen™ | Cat#AM2616 |
| Bovine Serum Albumin (BSA) | Jackson ImmunoResearch Labs | Cat# 001-000-162, RRID: AB_2336946 |
| Bovine Serum Albumin (IgG-Free, Protease-Free) | Jackson ImmunoResearch | Cat#001-000-162 |
| Normal Goat Serum | Jackson ImmunoResearch | Cat#005-000-121 |
| RNasin® Ribonuclease Inhibitor | Promega | Cat#N2515 |
| SUPERaseIn™ RNase Inhibitor | Thermo Fisher Scientific | Cat#AM2696 |
| Protease tablet | Roche | Cat#11873580001 |
| Laboratory stirrer | Yamato | Cat#LT-400 |
| IGEPAL CA-630 | Sigma-Aldrich | Cat#i8896 |
| Cell strainers | Falcon | Cat#352340 |
| Single cell partitioning | 10X Genomics | Chromium X |
| Chromium Next GEM Single Cell 3' Kit v3.1, 16 rxns | 10X Genomics | Cat#PN-1000268 |
| Streptavidin, Alexa Fluor™ 488 conjugate | Thermo Fisher Scientific | Cat# S32354 |
| Epredia™ 10% Neutral Buffered Formalin | Thermo Fisher Scientific | Cat#22-110-869 |
| Dynabeads™ MyOne™ Streptavidin T1 | Thermo Fisher Scientific | Cat#65602 |
| Phosphate-Buffered Saline (PBS) | Thermo Fisher Scientific | Cat#AM9625 |
| Hanks' Balanced Salt Solution (HBSS) | Thermo Fisher Scientific | Cat#14065056 |
| MgCl<sub>2</sub> (1M) | Thermo Fisher Scientific | Cat#AM9530G |
| KCl (2 M), RNase-free | Thermo Fisher Scientific | Cat#AM9640G |

### EXPERIMENTAL MODEL AND SUBJECT DETAILS

#### Mice

All experiments were approved by the Max Planck Florida Institute for Neuroscience Animal Care and Use Committee (Protocol #22-002) and were in accordance with the National Institute of Health Guide for the Care and Use of Laboratory Animals. Maximal efforts were made to reduce the suffering and the number of mice used. Male and female LepR-Cre (also named ObRb-Cre, Jackson Laboratory #008320), LSL-EGFP-L10 (Jackson Laboratory #030305), and C57BL/6J (Jackson Laboratory #000664) mice were 12–20 weeks old at the beginning of behavioral experiments. Animals (LepR-Cre and C57BL6J) were generally group housed, but kept in individual cages for free/operant feeding and drinking behaviors. Mice were kept under standard conditions in a day/night cycle of 12/12 hours (lights on at 7 am).

### METHOD DETAILS

#### Surgery and viral administration

Mice were anesthetized by isoflurane (3% for the induction and 1.5% during the surgery) and placed on a stereotaxic apparatus (Model 900, KOPF instruments, CA, USA) with a mouse adaptor and lateral ear bars. For viral vector delivery, AAV vectors were loaded in a glass pipette and infused into the brain by a nanoliter injector (DRUMMOND, nanoinject II). The coordinates in anteroposterior (AP) / mediolateral (ML) / dorsoventral (DV) from Bregma, were in mm: for viral injection to the insular cortex, +0.5/±3.85/-3.9 (250∼300 nL, 120 nL/min); for viral injection to the basal lateral amygdala -1.6/±3.3/-4.9 (150 nL, 120 nL/min); for cannula implantation to the insular cortex +0.5/±3.85/-3.6 (Plastics one, C315GMN/SPC); for canula connected with osmotic pump (ALZET, 2002) implantation to the insular cortex +0.5/±3.85/-3.9 (ALZET, Brain Infusion Kit 2, cat#0008663); for the ProView Integrated GRIN lens implantation to the insular cortex +0.5+3.85/-3.85 (Inscopix, 0.5mm × 6.1mm, 1050-004415); for the fiberoptic cannula implantation to the insula +0.5/±3.85/-3.8 (Doric Lenses, MFC_200/245-0.37_4mm_ZF1.25_FLT); for the fiberoptic cannula implantation to the basolateral amygdala -1.6/±3.3/-4.8 (Doric Lenses, MFC_200/245-0.37_6mm_ZF1.25_FLT). The coordinates used were decided according to the mouse brain atlas (Paxinos and Franklin, 2001, San Diego).

#### RNAscope (Fluorescence in situ hybridization technique)

C57BL/6J mice were euthanized by 5% isoflurane followed by cervical dislocation, after which brains were harvested and flash frozen on foil above dry ice. Samples were stored at -80°C before sectioning. Sections of 16 μm thickness were cut by cryostat (Leica CM3050 S) and stored in slide boxes at -80°C for up to three months. RNAScope Multiplex Fluorescent V2 Assay was then performed according to the manufacturer’s instructions using problems to visualize leptin receptor (ACDBio #402731) and VGAT (ACDBio #319191) mRNA, or leptin receptor and VGLUT1 (ACDBio #416631) mRNA. Briefly, sections were fixed for 15 minutes, then dehydrated with a series of ethanol washes in increasing concentration. A barrier was created around sections to contain solution with an ImmEdge hydrophobic barrier pen (Vector Labs, H4000), after which sections were incubated for 10 minutes in the HybEZ Humidity Control Tray (ACD, PN 310012) at room temperature. After washing and protease IV treatment, probes were added to each section and inserted into the HybEZ Oven (ACD, PN 321710) for 2 hours at 40°C. After washing, a series of amplification steps, HRP, and DAPI were added to the sections for visualization. Mounting medium (Fisher Scientific, 50-187-88) was then added to the slide and covered with a glass coverslip. Slides were dried overnight and imaged with Keyence BZ X810.

#### Histology

##### Nissl staining

Mice were perfused with a 10% formalin solution after sterile saline, their brains extracted and sectioned into 50 µm slices. Serial sections were made: one for sections that only received the Cre-dependent virus, and the other for sections that were to be Nissl stained. Sections for Nissl staining were mounted on a glass slide and dried for 1 hour. Sections were then demyelinated in an ascending graded ethanol series (70%, 95%, 100%) for 5 minutes each, and then rehydrated in a descending graded ethanol series (95%, 70%, 50%) for 5 minutes each. Following a rinse in milliQ water, shaken and filtered 1% cresyl violet solution was used to stain the sections, followed by rinsing in milliQ water three times. The sections were then destained in an ascending graded ethanol series (70%, 95%) for 5 minutes each and then for about 1 minute in a 100% solution. Finally, the sections were dried for 1 hour before being coverslipped with Fluoromount-G mounting medium. Brightfield images of the Nissl-stained sections were taken alongside fluorescent images of the mCherry-only sections using a Keyence fluorescent microscope (Keyenece BZ-X). Images of the Nissl-stained and fluorescent sections were paired up according to coronal depth and overlayed on Microsoft PowerPoint. Using the Allen Brain Atlas as reference, the cortical layers 1, 2/3, 4, 5, and 6 were drawn onto the Nissl-stained images with PowerPoint and then applied onto the fluorescent images (Figure 3). The number of the fluorescent cells in each cortical layer were quantified manually by a blinded experimenter.

##### Biocytin staining

Sections were placed in 4% paraformaldehyde (PFA) after at least 1 hour of whole-cell patch-clamp recording experiment (biocytin, Sigma-Aldrich cat#B4261, enters patched cells from internal solution of glass patch-clamp pipette), and were fixed over 48 hours. Sections were washed 3 times with PBS, then with 0.3% Triton X-100 in PBS (PBST). Streptavidin-488 (Thermo Fisher Scientific, S32354) was added with a 1:700 dilution in PBST. Sections were incubated overnight in the cold room (4°C) and then washed 3x with PBS. Sections were mounted, and images taken with a 40X water objective on a confocal microscope (Zeiss LSM 780). Images were analyzed by using Imaris 10.1.0.

##### mCherry staining

Similar to Nissl staining, LepR-Cre mice were injected with cre-dependent mCherry (pAAV-hSyn-DIO-mCherry, Addgene, 50459-AAV5, titer 2.3×10^13^ GC/mL) into the insular cortex. After 8 weeks mice were perfused with a 10% formalin solution after sterile saline, and had their brains extracted and sectioned into 50 µm slices from the frontal cortex to medulla. For the mCherry staining, the slices were washed three times with PBS for 5 minutes each while shaking, then once more with 0.3% PBST, then placed in 1 mL of blocking solution (0.3% PBST+3% BSA) while shaking at room temperature. The slices were then incubated with the primary anti-mCherry antibody (1:1000, abcam, ab205402, chicken polyclonal antibody) for around 48 hours at 4°C. The slices were moved out at room temperature for 1 hour, washed three times in PBS, and then incubated with the secondary goat anti-chicken-594 antibody (1:500, Thermo Fisher Scientific, A-11042) for 1 hour while shaking at room temperature. The slices were then washed three times in PBS, dried for 30 minutes, and coverslipped with Fluoromount-G with DAPI (Invitrogen, 00-495952) mounting medium. Images were taken with a Keyence fluorescent microscope (Keyence BZ-X800), and processed with Image J.

#### Electrophysiological patch-clamp recording

##### Brain tissue preparation

LepR-Cre mice were injected with cre-dependent mCherry (pAAV-hSyn-DIO-mCherry, Addgene, 50459-AAV5, titer 2.3×10^13^ GC/mL) or ChR2 (pAAV-EF1a-double floxed-hChR2(H134R)-mCherry-WPRE-HGHp, Addgene, 20297-AAV5, titer 2.4×10^13^ GC/mL) virus into the insular cortex. For mCherry expression, patch-clamp experiments were performed at least 1 week after surgeries. For ChR2 expression, patch-clamp experiments were performed at least 4 weeks after surgeries. Coronal brain slices (300 μm) were prepared in ice-cold cutting solution containing (in mM): 124 choline chloride, 26 NaHCO_3_, 2.5 KCl, 3.3 MgCl_2_, 1.2 NaH2PO_4_, 10 glucose and 0.5 CaCl_2_. After cutting, slices were allowed to recover for 30 minutes at 32°C and stored at room temperature in artificial cerebrospinal fluid (ACSF) containing (in mM): 124 NaCl, 26 NaHCO_3_, 3 KCl, 1.25 NaH_2_PO_4_, 20 glucose, 1 MgCl_2_, 2 CaCl_2_, 5 sodium ascorbate, 3 sodium pyruvate and 2 thiourea. All solutions were continuously bubbled with a carbogen mixture of 95% O_2_ and 5% CO_2_. Slices containing the insula or amygdala were transferred to a submersion recording chamber, superfused with oxygenated ACSF at a speed of 1–2 mL/minute.

##### Whole-cell patch-clamp recording for intrinsic excitability

Whole-cell patch-clamp recordings were performed using pipettes pulled from borosilicate glass capillaries (Sutter Instrument, BF150-110-10) with resistances of 3-4 MΩ. For recording intrinsic activity, a K-Gluconate based internal solution was used, containing (in mM): 85 K-Gluconate, 60 KCl, 5 NaCl, 10 HEPES, 0.5 EGTA, 4 MgATP, 0.3 Na_2_GTP, 5.37 Biocytin. An AMPA/kainite receptor antagonist, DNQX (20 μM), and 2 GABA receptor antagonists, picrotoxin (100 μM) and bicuculline (10 μM), were applied in the ACSF bath. LepR+ cells expressed mCherry in insula-claustrum region and were visualized by using epifluorescence microscope for whole-cell patch-clamp recording (Zeiss AXIO Eaxminer.D1). To measure the maximum firing rate, stimulation was gradually increased 20 times starting from -40 pA, with a step size of 20 pA. Each stimulation lasted for 1 second, with 10 seconds of interval between stimulations. The procedure was then repeated with 40 and 80 pA of step size. Leptin (working concentration: 100nM) was then placed into the bath, and the above stimulation procedure was then repeated after 15 minutes. Data were acquired with a Multiclamp 700B amplifier, Digidata1440 and Clampex software and analyzed with Clampfit software (Molecular Devices). Signals were filtered at 2 kHz and digitized at 10 kHz. Following patch-clam recording, patched sections were fixed by 4% PFA for at least two days, and then stained biocytin (filled to cell from internal solution in glass patch-clamp pipette, Sigma-Aldrich, B4261) and with streptavidine-488 (Thermo Fisher Scientific, S32354) to visualize the patched cell (see Biocytin staining method part for more details). The sections were imaged by using Zeiss LSM780, and images were analyzed by using Imaris 10.1.0.

##### Whole-cell patch-clamp recording for paired-pulse ratio and AMPA/NMDA ratio

Whole-cell patch-clamp recordings were performed using pipettes pulled from borosilicate glass capillaries (Sutter Instrument, BF150-110-10) with resistances of 3-4 MΩ. For recording postsynaptic currents, we used Cs-Gluconate based internal solution containing (in mM): 140 Cs-Gluconate, 8 NaCl, 10 HEPES, 0.2 EGTA, 2 MgATP, and 0.2 Na_2_GTP. LepR+ cells expressed ChR2 in the insula-claustrum region. ChR2 labeled terminals in insula or basolateral amygdala were stimulated with 470-nm light pulses (1–5 ms) from a light-emitting diode (pE-100, Cool LED LTD) through the 20X (4X optical zoom) 1.0 NA objective (Zeiss AXIO Examiner.D1) or through the 40X 1.0 NA objective (Olympus, BXWI-5). For demonstration of monosynaptic inputs, TTX (1uM) and 4-AP (1mM) were applied in the ACSF bath. The Paired Pulse Ratio is the ratio of the amplitude of the second response to that of the first where 2 sequential light pulses are delivered with different apart, 25, 50, 100 or 200 ms. Data were acquired with a Multiclamp 700B amplifier, Digidata1440 and Clampex software and analyzed with Clampfit software (Molecular Devices). Signals were filtered at 2 kHz and digitized at 10 kHz. For analysis of AMPA / NMDA ratio, the AMPA component was isolated at -70 mV where NMDA receptors are blocked by Mg^2+^ and in presence of 100 µM Picrotoxin to block GABA_A_ currents. The NMDA current was measured at +40 mV, 50ms after the start of synaptic response such that the AMPA current with fast decay kinetics is negligible. NMDA currents were blocked by 50uM AP5. Further analysis and graphs were completed in Microsoft Excel and GraphPad Prism 8.0.

#### Leptin infusion

Leptin (R&D, 498-OB-1M) stock solution was stored in -80°C freezer and was used within 3 months after the stock preparation. In the acute leptin administration experiment, the leptin working solution was 400ng/μL., and 0.3 μL of the working solution was injected into the insula by micropump (Harvard Apparatus, Pump 11 Picro Plus Elite) with a speed of 0.1 μL/minute. Food intake and water intake were monitored daily in the home cage, and was also measured at 1, 2, 4, 12, 24 hours after acute leptin infusion. Food intake was measured by a food dispenser (FED 3.0^40^, using Dustless Precision Pellets® Rodent, Grain-Base #F163). Water intake was measured using water bottles assembled by Nalgene™ PPCO Conical-Bottom Centrifuge Tubes (Thermo Fisher Scientific, 3103-0050PK), a rubber stopper (Eisco, CH0314K1H), and drinking tube (Ancare, TD100). In the chronic leptin administration experiment, the leptin working solution was 10 ng/μL, and around 200 μL of working solution was loaded to the osmotic pump; the delivery speed to the insula was 0.5 μL/hour and the delivery lasts for up to 14 days. Food intake and water intake were monitored daily in the home cage and normalized to baseline. The baseline of food intake and body weight was daily measured for a week before osmotic pump implantation. For feeding measurement after food restriction, mice were changed to clean cages prior to an 18-hour food restriction. Leptin infusion (120 ng) was performed either before food restriction or prior to refeeding. Food intake was recorded by FED 3.0 and body weight was measured daily. For the progressive ratio test, mice were under food restriction in which 2∼2.5 grams of standard 5V5R chow were provided daily for a week and the body weight reached to 85-90% of body weight prior to food restriction. Mice were training on a fixed ratio 1 (FR1) schedule (Pellet delivery requires one-time nosepoke) for 3 sessions (60 minutes/session, a session daily), and fixed ratio 5 (FR5, 5 nosepokes for one pellet) for 3 sessions. Finally, mice were tested In the progressive ratio task, where the required nosepoke number increases with each pellet retrieval according to the following schedule: ((5 * exp ((Pellet number + 1) * 0.2)) - 5). Mice were under food restriction during training and testing of progressive ratio, but were sated at the test given 6 days after leptin infusion.

#### Viral-TRAP

Affinity purification of EGFP-tagged polysomes was conducted in LepR-Cre mice crossed with LSL-EGFPL10 to tag LepR+ polysomes. Polysomes were immunoprecipitated according to the protocol in Nectow et al., 2017^44^. Briefly, mice were separated into three biological replicate groups of 4–6 mice per group and euthanized. Brains were extracted, and the insular cortices were dissected on ice and pooled. Tissue was homogenized in buffer containing 10 mM HEPES-KOH (pH 7.4), 150 mM KCl, 5 mM MgCl_2_, 0.5 mM DTT, 100 μg/mL cycloheximide, RNasin (Promega, Madison, WI, 2515) and SUPERase-In (Life Technologies, Waltham, MA, AM2696) RNase inhibitors, and complete-EDTA-free protease inhibitors (Roche, 4693124001) and then cleared by two-step centrifugation to isolate polysome-containing cytoplasmic supernatant. Polysomes were immunoprecipitated using monoclonal anti-EGFP antibodies (clones 19C8 and 19F7; see Heiman et al., 2008) bound to biotinylated-Protein L (Pierce; Thermo Fisher Scientific, 29997)-coated streptavidin-conjugated magnetic beads (Thermo Fisher Scientific, 65602). A small amount of tissue RNA was saved before the immunoprecipitation (Input) and both input and immunoprecipitated RNA (IP) were then purified using RNAeasy Mini kit (QIAGEN, 74104). RNA quality was checked using an RNA PicoChip on a Bioanalyzer (Agilent 2100). RIN values > 7 were used. Experiments were performed in triplicates for each group. cDNA libraries were prepared using NEBNext Ultra II DNA Library Prep Kit and sequenced on an Illumina NextSeq 2000 platform with 2X50bp P3 sequencing.

##### qPCR for Viral-TRAP validation

cDNA was prepared with the QuantiTect Reverse Transcription Kit (QIAGEN). qPCR using predesigned Taqman probes (idtDNA) were used. The abundance of these genes in IP and Input RNA was quantified using Taqman Gene Expression Master Mix (Applied Biosystems #4444556) and run on a Applied Biosystems QuantStudio 7. Transcript abundance was normalized to beta-actin. Fold change was calculated using standards or with the ΔΔCt method if there was not enough material to make standards.

#### 10X single cell sequencing

##### Tissue processing and isolation of nuclei

Mice were treated by i.p. DPBS or Leptin (5 mg/kg of body weight). 24 hours after the treatment, mice were euthanized by 5% isoflurane followed by cervical dislocation. After dislocation, brains were rapidly dissected and placed in an adult mouse brain slicer matrix on ice, and cut to 1.0 mm thick coronal slices which were obtained by inserting 4-5 GEM blades (Electron Microscopy Sciences, 71970-WA) in the matrix. All the following steps were performed strictly on ice. Bilateral sections of insular cortex tissue were dissected from the tissue punches, then transferred to pre-cooled 10 mL tubes on ice and homogenized with pestles 10 times in 0.5 mL of ice-cold Complete HB Buffer, which was prepared by 3 mL Buffer HB (mixed 50 mL 0.5 M sucrose, 2.5 mL 1 M KCl, 0.5 mL 1 M MgCl_2_, 2.66 mL 0.75 Tricine-KOH, pH 7.8, and 44.34 mL H_2_O), 4.5 μL RNasin Plus RNase inhibitor (Promega, N2515), 3.6 μL 35% BSA (Invitrogen, AM2616), 3 μL spermine (Sigma, S1141), 3 μL spermidine (Sigma, S2501), 3 μL DTT (Sigma, D0632) and one third of Protease tablet (Roche, 11873580001). The tissue was homogenized with a tight pestle for 10 times by using a laboratory stirrer (Yamato LT-400) with a speed around 800 rpm. Then, 16 μL of 5% IGEPAL CA-630 solution (Sigma, I8896) was added and the tissue was immediately homogenized 5 more times. The cell suspensions were then passed through 40 μm cell strainers (Falcon, 352340) into 2 mL conical tubes. The filtered homogenate of each sample was combined with 0.5 mL of Working Solution, which was prepared by mixed 2.5 mL Optiprep, 0.5 mL Diluent (150 mM KCl, 30 mM MgCl_2_, 120 mM Tricine-KOK, pH 7.8), 3.6 μL BSA and 4.8 μL RNAsin, before layering on top of 30% (mixed 0.45 mL working solution and 0.3 complete HB buffer) and 40% (mixed 0.45 mL working solution and 0.3 complete HB buffer) iodixanol gradient solutions. Cells were centrifuged for 40 minutes at 4,500 × g speed at 4°C with full acceleration and braking. After centrifugation, 60 μL of each nuclei sample was collected at the 30%-40% iodixanol interface. 6 μL of nuclei were collected from each sample and mixed with 6 μL of Trypan blue (Gibco, 15-250-061) for manual cell counting and nuclei quality assessment. Nuclei were counted in a hemocytometer (Bright-Line, Z359629-1EA) using a brightfield microscope (OLYMPUS, CKX41). Working solution, Complete Buffer HB, 5% IGEPAL CA-630, and iodixanol solutions were freshly prepared the day of the experiment. Stock buffers of Diluent, Buffer HB, spermine, spermidine, and DTT were prepared less than one month before use and stored in the -20°C freezer. Stock buffers were thawed on ice before tissue collection.

##### cDNA library construction and sequencing

Following nuclei isolation, approximately ten thousand nuclei from each sample were subjected to the Chromium Next GEM Single Cell 3’ Gene Expression protocol (10X Genomics) according to the manufacturer’s instructions. cDNA libraries were constructed by using the protocol and fragment sizes were quantified by Tapestation (Agilent, 4150) for quality control analysis and sequenced (2×300 bp, paired-end reads configuration) using the NextSeq 2000 (Illumina). Final 10X snRNA-seq cDNA Libraries were sequenced at a minimum of 20,000 read pairs/cell. **DPBS and leptin groups were sequenced separately on an Illumina NextSeq 2000 apparatus.**

#### *In vivo* calcium imaging

LepR-Cre Mice were injected with pGP-AAV-syn-FLEX-jGCaMP8m-WPRE (Addgene, 162378-AAV1, 1.7×10^13^ GC/mL) into the insula cortex, followed by GRIN Lens implantation. C57BL/6J Mice were injected pGP-AAV-syn-jGCaMP8m-WPRE (Addgene, 162378-AAV1, 2.5×10^13^ GC/mL) into the insula cortex, followed by GRIN Lens (Inscopix, ProView^TM^ Integrated Lens 0.5mm × 6.1mm, 1050-004415) implantation. Animal behavior commenced 6 weeks after surgeries. Mice were first habituated to the miniscope mounted before imaging began.

1) <u>Food restriction:</u> Adult mice over 20 grams received ∼70% (2∼2.5 grams of standard 5V5R chow food) of daily food intake before food restriction. Food was placed on the floor of home cages. After ∼1 week of food restriction, body weight reached to 85-90% of body weight after which training was begun.
2) <u>Nose poke feeding</u>: Using a FED 3.0 device, mice were required to nosepoke in order to get an available food pellet. Nosepoke ports were available both on the left and right of the food well. Two seconds after a nosepoke followed by a 0.2-second sound from a buzzer and 0.2-second blue light in FED 3.0, a pellet (Dustless Precision Pellets® Rodent, Grain-Base #F163) was delivered to the pellet well. The response period was 8 seconds after the nosepoke, during which mice could not obtain a second pellet. Mice were provided extra chow to reach 70% (∼2g) of daily food intake if mice ate less than 2g food during the behavioral task. This training phase lasted for 8-10 sessions (one 30-minute session per day) and mice could obtain 20 pellets in a session. Miniscope (Inscopix nVue) imaging began after the training phase, with a habituation session (mounted miniscope without imaging). Imaging experiments were performed the day after habituation, and imaging lasted for the entire 30-minute behavior session.
3) <u>Nose poke feeding and drinking choice</u>: Two seconds after a nose poke followed by a 0.2- second sound from a buzzer and 0.2-second blue light in FED 3.0, a pellet was delivered to the pellet well and the water spout became active. Mice could choose to get 5µl water by licking the active water spout 2 seconds after nose poke or to retrieve the food pellet. If mice obtained the pellet first, the water spout became inactive. If mice licked water first, the pellet was still available in the well. Thus, lick-then-feed options were available, but not feed-then-lick. The training phase lasted for 3∼7 sessions under food restriction conditions, until mice obtained 5 water rewards, after which the imaging session was conducted. Following imaging under food restriction, mice then were switched to water restriction and performed the same behavior for 2 sessions. Mice were provided extra water to reach to 1 mL and 0.8 mL of consumed water for the first day and second day after training, respectively, after which the second imaging session was conducted.
4) <u>Free feeding and drinking behaviors</u>: Mice were able to freely obtain food and water freely in the behavioral chamber for 3-5 sessions (one session per day, each session lasted 30 minutes) under food restriction, after which one 20-30 minutes session was conducted in conjunction with calcium imaging. Mice were provided extra chow to reach to ∼2g food if mice ate less than 2g of food pellet from FED3.0 feeder during the behavioral task. Mice were then switched to water restriction for two days (provided 1 mL and 0.8 mL of water daily for the first day and second day), after which imaging was conducted again for one session.

#### Retrograde viral tracing for Cre+ and Cre-cells

A retrograde color-switching virus (100 nL, pOTTC1032-pAAV EF1a Nuc-flox(mCherry)-EGFP, Addgene, Addgene, 112677-AAVrg, titer 2.5×10^13^ GC/mL) was injected into the basolateral amygdala of LepR-Cre mice. The virus expresses EGFP in Cre positive (Cre+) cells, and mCherry in Cre negative (Cre-) cells. Mice were perfused with 10% formalin solution 4 weeks after viral injections. Brains were sectioned on a vibratome (Leica VT1000S) at 50 μm sections and mounted on slides with DAPI, as described above. Images were taken by using a Keyence BX810 microscope.

#### Simultaneous whole-cell patch-clamping and calcium imaging in vitro

LepR-Cre mice were injected with mixture of 50% Cre-dependent mCherry (pAAV-hSyn-DIO-mCherry, Addgene, 50459-AAV5, titer 2.3×10^13^ GC/mL) and 50% pGP-AAV-syn-jGCaMP8m-WPRE (Addgene, 162378-AAV1, 1.7×10^13^ GC/mL) into the insula cortex. Whole-cell patch-clamping of LepR+ cells in the insula was as the above-mentioned method and simultaneously performed with calcium imaging (Zeiss AXIO Eaxminer.D1). Imaging had a 20 Hz acquisition speed. First, 175 frames of GCaMP images were taken in a 100×100 µm field, followed by 1 second of current injection (50 mV, voltage clamp recording) to the patched LepR+ cell, and then 250 additional frames of CCaMP images. This procedure was repeated 3 times. The images were analyzed by using ImageJ.

#### Optogenetic stimulation during operant behavior

LepR-Cre mice were injected with ChR2 virus (pAAV-EF1a-double floxed-hChR2(H134R)-mCherry-WPRE-HGHpA, 2.4×10^13^ GC/mL, Addgene 20297-AAV5) or mCherry virus (pAAV-hSyn-DIO-mCherry, 2.3×10^13^ GC/mL, Addgene 50459-AAV5) into the insular cortex. A fiberoptic was then implanted to the insula or the basolateral amygdala. Mice were trained for nose poke feeding as described above, under food restriction. For the stimulation experiments, mice were divided to two groups. In the first day, one group received stimulation, while the other group of mice were connected to the patch cord without light on. The next day the groups were reversed and the protocol repeated. Two stimulation protocols were used: 1) Stimulation for 10 minutes before providing pellets after nose poke. Stimulation was given with blue light on every other second (20 Hz, 5 ms/pulse) by using 470 nm LED (ThorLabs, M470F4). After 10 minutes, operant feeding behavior began and lasted for 20 minutes. 2) Stimulation after nose poke. There was a 50% probability of stimulation (blue light 20 Hz, 5 ms/pulse) after nose poke and the stimulation lasted for 2 seconds. Stimulation experiments were performed for 2∼3 sessions and there were at least two days between protocol 1 and 2.

#### Real time place preference paradigm (RTPP)

RTPP was modified from a previous protocol^66^. The experiment was performed using a three-compartment chamber, including two main compartments (40 cm (W) × 20 cm(L) × 20 cm(H)) and small connecting compartment (20 cm (W) × 20 cm(L) × 20 cm(H)). Mice were habituated for 15 minutes without connecting patch cord. The next day, mice were given a pretest of 15 minutes with the patch cord connected and initial preferences were recorded. On the third and fourth day, RTPP was tested 30 minutes, in which stimulation was given when mice entered the right-side compartment (light on every other second with blue light (20 Hz, 5 ms/pulse) by using 470 nm LED). The next day we tested the conditioned response (CR) by placing mice into the chamber for 15 minutes without connecting the patch cord; after 2∼4 days before next step, we conducted days 3-5 but reversed the side of stimulation.

### QUANTIFICATION AND STATISTICAL ANALYSIS DATA ANALYSIS

All results are presented as mean ± SEM. and were analyzed with Prism software or Matlab. No statistical methods were used to predetermine sample sizes, but our sample sizes are similar to those reported in previous publications. Normality tests and F tests for equality of variance were performed before choosing the statistical test. Unless otherwise indicated, statistics were based on unpaired-t tests, Wilcoxon matched-pairs signed rank test, one-way ANOVAs or two-way ANOVAs with Sidak’s posthoc comparison. p < 0.05 was considered significant (∗p < 0.05, ∗∗p < 0.01, ∗∗∗p < 0.001, ∗∗∗∗p < 0.0001). Animals in the same litter were randomly assigned to different treatment groups and blinded to experimenters in the various experiments. Injection sites and viral expression were confirmed for all animals. Mice showing incorrect injection sites or optic fiber placement were excluded from the data analysis.

#### snRNA sequencing data primary processing and visualization

Analysis was performed at The Herbert Wertheim UF Scripps Institute for Biomedical Innovation & Technology, Bioinformatics and Statistics Core Facility (RRID:SCR_023048). The quality of the reads remained high throughout the sequencing process. The reads were tested for the presence of rRNA, tRNA, mitochondrial, and mycoplasma sequences, common contaminants in RNA-seq experiments. Each sample contained less than 1% rRNA and no mycoplasma. Individual samples showed no significant differences in terms of the number of reads and the proportion of reads that were either trimmed or unable to be mapped to the genome. The RNA sequencing pipeline consisted of Python version 3.7.3, Cutadapt version 3.4, STAR aligner version 2.7.5c, HTSeq version 0.11.2, R version 4.2.3, DESeq2 version 1.42.1. Genome Mapping was performed based on the mouse genome (mouse-ENSEMBL-GRCm38.r91: M.musculus-ENSEMBL-GRCm38.r91; downloaded February 16, 2018). DESeq2 package (default settings) was used to identify differentially expressed genes (FDR<0.05), and PCA was depicted to detect outliers and batch effects using the 500 most variable genes.

Raw base call (BCL) files were analyzed using CellRanger (v7.0.0) (PMID: 28091601). The “mkfastq” command was used to generate FASTQ files and the “count” command was used to generate raw gene – cell expression matrices. Ambient RNA contamination was inferred and removed using CellBender (v0.2.0) with standard parameters. Nuclear fraction that quantifies the amount of RNA in a droplet that originated from unspliced pre-mRNA were calculated for every cell barcode in the sample BAM file using DropletQC (v.0.0.0.9000)^67^. Mouse genome mm10 was used for the alignment and gencode vM25 was used for gene annotation and coordinates^68^. Downstream analysis was performed in R with Seurat (v4.3.0)^69^ and customized R scripts. Data from different samples (Leptin and DPBS) were merged into a unique single cell object. Matching each cell bar code, nuclear fraction score was included in the metadata. Filtering was conducted by retaining nuclei that had unique molecular identifiers (UMIs) less than 15000, mitochondrial content less than 5 percent, number of Genes (nGene) greater than 250 and nuclear fraction (nf) greater than 0.5. Doublets were removed using scDblFinder ((v1.12.0)^70^ for a resultant expression matrix with 21,849 genes and 119,694 cells. The SCTransform workflow was used for count normalization, initial integration, and to identify highly variable genes^71^ using 30 principal components and resolution of 0.5 for Louvain clustering and UMAP. Batch correction was performed using Harmony (v0.1.1)^72^. Cluster marker genes were identified using FindAllMarkers using Wilcoxon Rank Sum test with the standard parameters. We identified 26 final distinct clusters, of which two clusters showing high expression of Striatum were removed. Cell annotation was done by manual curation of known canonical markers. Pseudobulk differential expression analysis: To identify differentially expressed genes between conditions (Leptin vs Saline) in each defined cell types was found with the R package DESeq2 (v1.44.0)^73^. Genes were defined as significantly differentially expressed at padj < 0.05 and abs (log2FoldChange) > 0.3. Gene ontology analyses: The functional annotation of the identified DEGs was performed using scToppR (v0.99.1)^52^. Enrichment was further confirmed using the R package cluster Profiler (v4.12.2)^74^. Cell Perturbation Analysis: Cell perturbation analysis was conducted using MELD (v1.0.2)^50^. A MELD model with default parameters was fit to the entire dataset, using the ‘Exp_Grp’ (Leptin-treated vs. DPBS control) as the dependent variable, calculating the distribution of likelihoods for different cell types being most altered by the Leptin treatment. Augur (v1.0.3)^51^ followed by a logistic regression as classifier was used to further confirm the magnitude of leptin-induced response of each cell-type. Data availability: Raw and processed data to support the findings of this study have been deposited in GEO under accession number: TBD. All other data supporting the findings of this study are available from the corresponding author on reasonable request. Source data are provided with this paper. Code availability: Custom code used is available from GitHub at https://github.com/BioinformaticsMUSC/SternLab_Leptin_Codes Interactive web app is available at https://bioinformatics-musc.shinyapps.io/SternLab_Leptin_Study/Acknowledgements.

#### *In vivo* calcium imaging and behavioral data

Behavioral data was collected by FED 3.0 and MATLAB. Calcium imaging data was collected by Inscopix DAQ box with 20 Hz acquisition speed and preprocessed by Inscopix IDPS software. Raw imaging data was reduced to 10 Hz from 20 Hz, then ΔF/F values of individual cells were obtained using the PCA/ICA approach. Further data analysis was done using custom codes in MATLAB. The individual cell ΔF/F values were normalized to a range of 0 to 1 by the function, ΔF/F normalization = (Data - Data_Minimum_) / (Data_Maximum_ – Data_Minimum_). For the heatmap graphs and hierarchical cluster analysis, the variance of data (z-score) was applied.

#### K-Nearest Neighbors (KNN) and Gaussian Naïve Bayes

To classify population neural activity for decoding animal behavior, cell activity time series were first aligned with the behavior label with predefined short time window [-5s, 5s] (pre-event, post-event). Two classifiers, KNN and Gaussian Naive Bayes, were used to evaluate their ability to distinguish different behaviors, including nose poke and feeding. This evaluation was carried out using repeated stratified 4-fold cross-validation, ensuring that each fold maintained a balanced representation of the classes. For each repetition and fold, the models were trained and tested with both overall and per-class accuracies computed. To further validate the significance of the classification results, a control analysis was performed in which the labels were randomly shuffled, providing a baseline to assess whether the observed performance is meaningful.

#### Hierarchical clustering analysis

To group cells based on their activity patterns, hierarchical clustering was performed. The dataset used was the average activity of cell traces aligned at the event onset, where each row was an average cell trace. A linkage matrix was computed from the cell activity time series using the ’complete’ linkage method, which creates a tree of hierarchical clusters of the rows of the dataset based on the Euclidean distances between rows. The resulting hierarchical tree was then segmented into four clusters calculated by Calinski-Harabasz criterion, assigning each cell to one of the four groups based on similarity in activity profiles.

To quantify the temporal dynamics in neural population activity, we computed the autocorrelation half-width (ACHW) for each cell’s activity time series. ACHW reflects how long the temporal structure of a cell’s activity remains correlated with itself, serving as a proxy for the temporal persistence or rhythmicity of neural responses under different behavioral conditions. For each cell’s activity trace, the normalized autocorrelation was calculated. The ACHW was defined as the first positive time lag at which the autocorrelation dropped below 0.5. This analysis was applied to cell activity recorded during different behavioral conditions. ACHW values were compared across conditions using a Kruskal-Wallis test, followed by multiple comparisons.

#### Principal component analysis (PCA)

To reduce dimensionality and visualize the variance in cell activity, principal component analysis (PCA) was applied. The dataset was same as the one for above hierarchical cluster analysis. Prior to PCA, the data were z-scored across time for each cell to normalize activity aligned at the event onset. PCA uses the singular value decomposition algorithm. The first three principal components were extracted and visualized in a 3D plot, where each point represents population activity at a timepoint in the reduced feature space. Color coding was used to indicate temporal conditions from 5 s prior to the event onset and 20 s posterior to the event.

#### Dissimilarity analysis

To assess the similarity of cell activity between feeding and drinking conditions, principal component analysis (PCA) was first performed separately on each dataset. The first three principal components from each condition were then compared using two complementary approaches:

1. Procrustes analysis was used to calculate the geometric similarity between the PCA spaces by aligning the first three principal components from each condition. Procrustes determines a linear transformation (translation, reflection, orthogonal rotation, and scaling) of two trajectories in PCA spaces. The resulting Procrustes distance reports the minimum value of dissimilarity. Smaller values indicate greater similarity in the spatial structure of the components.
2. Subspace analysis was used to calculate the principal angle between two trajectories in 3D PCA subspaces. First, the orthonormal bases were computed by using singular value decomposition in “orth”. Then the angle of two projections was computed. A smaller angle (i.e., larger cosine similarity or singular values) suggests a higher degree of alignment between the underlying activity patterns in the two conditions.

