## Supplementary Figures for "Direct interoceptive input to the insular cortex shapes learned feeding behavior"

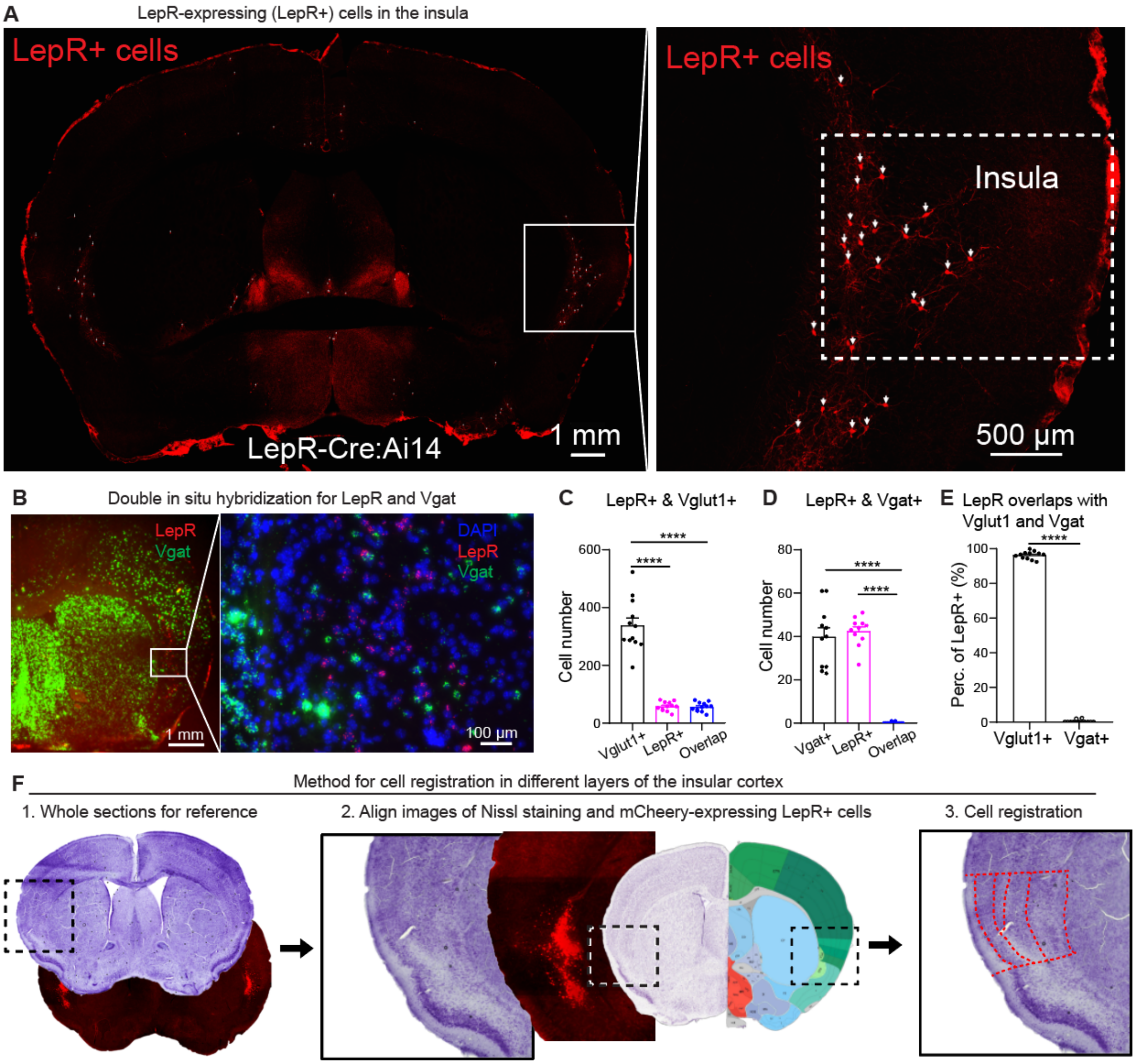


**Figure S1.** Characterization of $INS^{LepR}$ cells in cell types and cortical layers, related to **Figure 1**.

(A) TdTomato (Ai14), expressed in LepR+ cells in the insular cortex. White arrows point to TdTomato+ cells. The inset on the left is magnified on the right.
(B) Two-color fluorescent *in situ* hybridization for LepR mRNA (red) and Vgat mRNA (green). The inset on the left is magnified on the right.
(C) Quantification of the number of cells co-expressing (blue) LepR mRNA (magenta) and Vglut1 mRNA (black) (n = 12 in each group, one-way ANOVA followed by Tukey post-hoc test, $F_{(2,33)}$=110.4, **** $p$ < 0.0001; Vglut1+ vs. LepR+, **** $p$ < 0.0001; Vglut1+ vs. Overlap, **** $p$ < 0.0001).
(D) ) Quantification of the number of cells co-expressing (blue) LepR mRNA (magenta) and Vgat mRNA (black) (n = 11 in each group, one-way ANOVA followed by Tukey post-

hoc test, $F_{(2,30)}$=77.5, **** $p < 0.0001$; Vgat+ vs. Overlap, **** $p < 0.0001$; LepR+ vs. Overlap, **** $p < 0.0001$).

(E) Fraction of LepR+ cells, expressed as a percentage (%), that coexpress either Vglut1+ or Vgat+ (n = 12 Vglut1+, n = 11 Vgat+, two-tailed unpaired t test, **** $p < 0.0001$).

(F) Method for registration of LepR+ cells expressing mCherry in different layers of the insular cortex to Nissl staining.

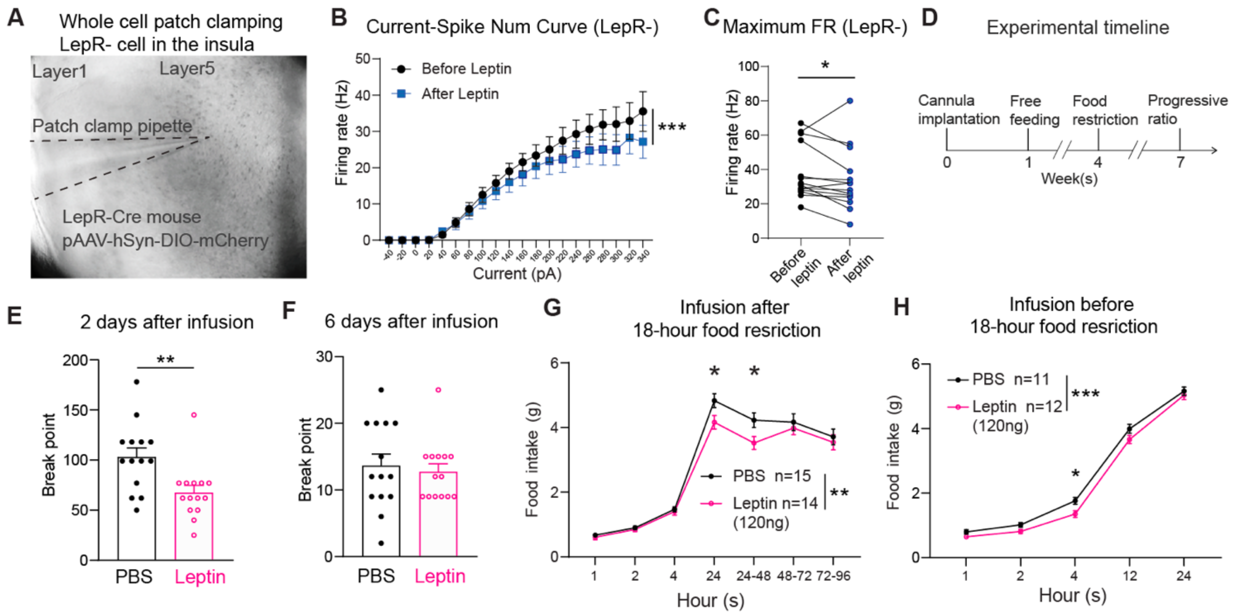


**Figure S2**. Leptin effects on intrinsic excitability and feeding, related to **Figure 1**.

(A) Image of a representative brain section with patch clamp pipette for recording of LepR- cells in the insular cortex.
(B) Firing rates in LepR- cells triggered by a series of currents before (black) and after (blue) leptin administration (n = 7-14 cells in each group from 6 mice, two-way ANOVA followed by Tukey post-hoc test, $F_{(1,460)}$ = 12.36, *** *p* = 0.0005).
(C) Maximum firing rate (FR) in LepR- cells triggered by current injection before and after leptin administration (n = 14 cells from 6 mice, Wilcoxon matched-pairs signed rank test, * *p* = 0.0289).
(D) Overall experimental timeline for experiments with local injection of PBS (control) or leptin.
(E) Breakpoint in the progressive ratio task, tested 2 days after PBS or leptin infusion under food restriction (n = 14 PBS, n = 14 leptin, two-tailed unpaired t test, ** *p* = 0.0053).
(F) Breakpoint in the progressive ratio task, tested 6 days after PBS or leptin infusion under food ad libitum condition (n = 14 PBS, n = 14 leptin, two-tailed unpaired t test, *p* = 0.6691).
(G) Food intake measurements, expressed in grams (g), following an 18-hour food restriction. PBS or leptin (120ng) was infused into the insula immediately following the 18-hour food restriction (n = 14-15 PBS, n = 14-15 leptin, two-way ANOVA followed by Tukey post-hoc test, $F_{(1,187)}$ = 8.465, ** *p* = 0.0041. PBS vs leptin, at 24-hour, * *p* = 0.0497; PBS vs leptin, at 24-48 hours, * *p* < 0.0330).
(H) Food intake measurements, expressed in grams (g), following an 18-hour food restriction. PBS or leptin (120ng) was infused into the insula before 18-hour food restriction (n = 11 PBS, n = 12 leptin, two-way ANOVA followed by Tukey post-hoc test, $F_{(1,105)}$ = 13.74, *** *p* = 0.0003. PBS vs leptin, at 4-hour, * *p* = 0.0310).

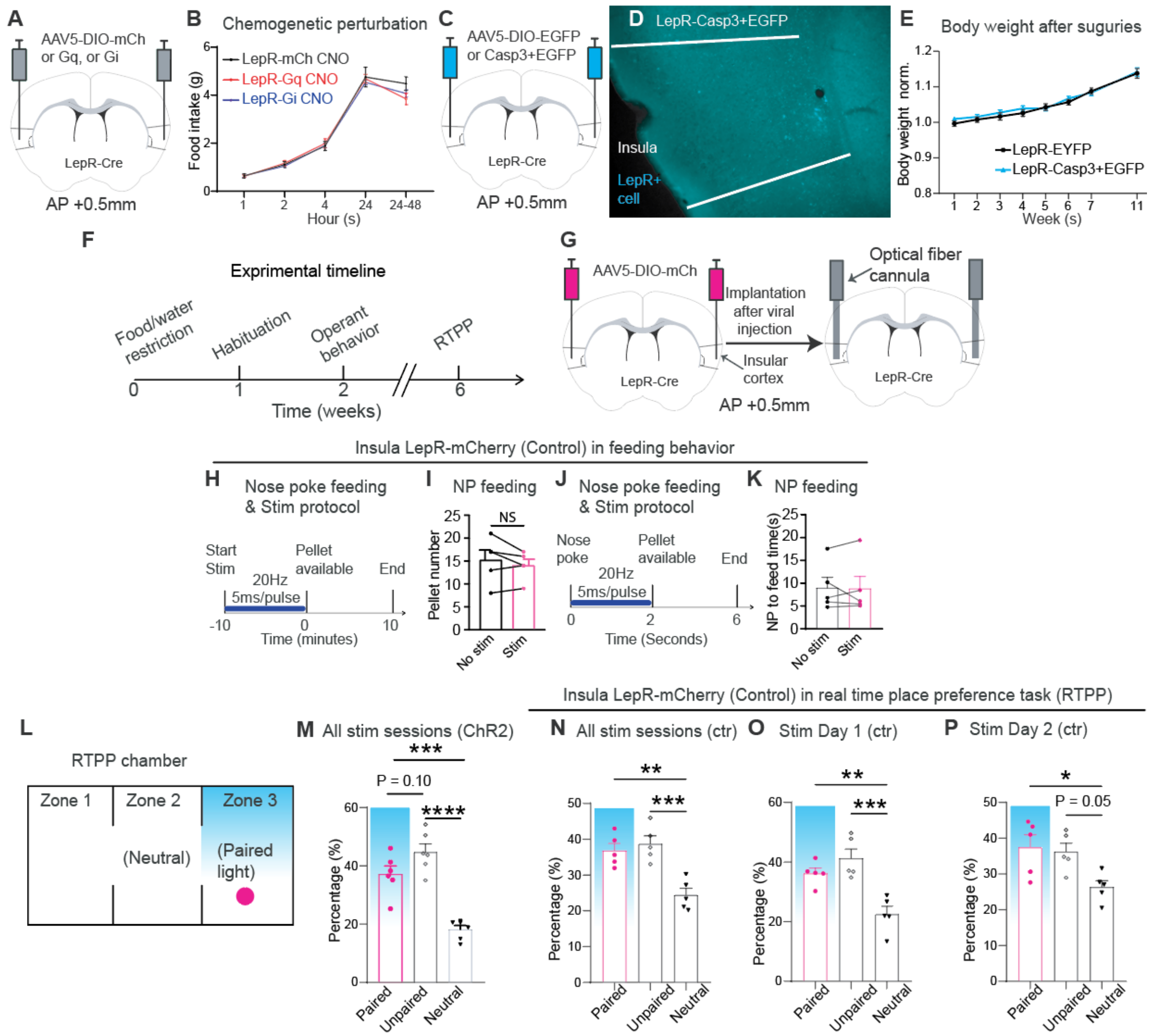


**Figure S3.** Chemogenetic and optogenetic control of INS$^{LepR}$ and effects on feeding and place preference behavior, related to **Figure 2**.

(A) Scheme for the expression of DREADD-Gq or Gi or mCherry control (mCh) in INS$^{LepR}$ cells.
(B) Food intake measurements, expressed in grams (g), following chemogenetic activation (Gq, red) or inhibition (Gi, blue) of INS$^{LepR}$ cells. Food intake measurement started 30 minutes after i.p. CNO (2 mg/kg) injection.
(C) Scheme for the expression of Caspase3 or control (EGFP) in the INS$^{LepR}$ cells.
(D) Image showed ablation of LepR+ cells (cyan) following Caspase3 expression.
(E) Normalized body weight following Caspase3 expression in the INS$^{LepR}$ cells.
(F) Scheme for the experimental timeline of behaviors for optogenetic experiments.
(G-K) Behavioral results for control experiment with optogenetic stimulation in mCherry-injected mice.

(G) Scheme for the expression of mCherry control virus in $INS^{LepR}$ cells and optical fiber cannula implantation.
(H) Scheme for the protocol of pre-session stimulation (blue).
(I) Pellet number consumed over one testing session with pre-session stimulation (magenta) or control (blue) (n = 5 Stim, n = 5 No stim, two-tailed paired t test, $p$ = 0.3046).
(J) Scheme for the protocol of post-nosepoke stimulation. Stimulation is given for 2 seconds following the nosepoke before food and water choice is available.
(K) (H) Latency to pellet retrieval following nose poke with stimulation (magenta) or control (blue) (n = 6 Stim, n = 6 No stim, two-tailed paired t test, * p = 0.0712). (n = 5 Stim, n = 5 No stim, two-tailed paired t test, $p$ = 0.8884).
(L) Scheme for the real-time place preference (RTPP) chamber coupled with optogenetic stimulation. Stimulation is depicted in Zone 3, but switches between Zone 1 (Days 3 and 4) and Zone 3 (Days 6-7).
(M) Percentage of time staying in the stimulation paired zones, averaged across all of the stimulation sessions (n = 6, one-way ANOVA followed by Tukey post-hoc test, $F_{(2,15)}$ = 31.49, **** $p$ < 0.0001; Paired vs. Unpaired, $p$ = 0.1017; Paired vs. Neutral, *** $p$ = 0.0002; Unpaired vs. Neutral, **** $p$ < 0.0001).
(N-P) RTPP test in control mice with expression of mCherry in $INS^{LepR}$ cells.
(N) Percentage of time staying in Zone 1-3 on Stim Day 1 and 2 with insula LepR-mCherry mice (n = 5, one-way ANOVA followed by Tukey post-hoc test, $F_{(2,12)}$ = 14.35, *** $p$ = 0.0007; Paired vs. Unpaired, $p$ = 0.7929; Paired vs. Neutral, ** $p$ = 0.0028; Unpaired vs. Neutral, *** $p$ = 0.0009).
(O) Percentage of time staying in Zone 1-3 on Stim Day 1 with insula LepR-mCherry mice (n = 5, one-way ANOVA followed by Tukey post-hoc test, $F_{(2,12)}$ = 14.16, *** $p$ =0.0007; Paired vs. Unpaired, $p$ = 0.3799; Paired vs. Neutral, ** $p$ = 0.0071; Unpaired vs. Neutral, *** $p$ = 0.0007).
(P) Percentage of time staying in Zone 1-3 on Stim Day 2 with insula LepR-mCherry mice (n = 5, one-way ANOVA followed by Tukey post-hoc test, $F_{(2,12)}$ = 5.244, * $p$ = 0.0231; Paired vs. Unpaired, $p$ = 0.9423; Paired vs. Neutral, * $p$ = 0.0301; Unpaired vs. Neutral, $p$ = 0.0538).

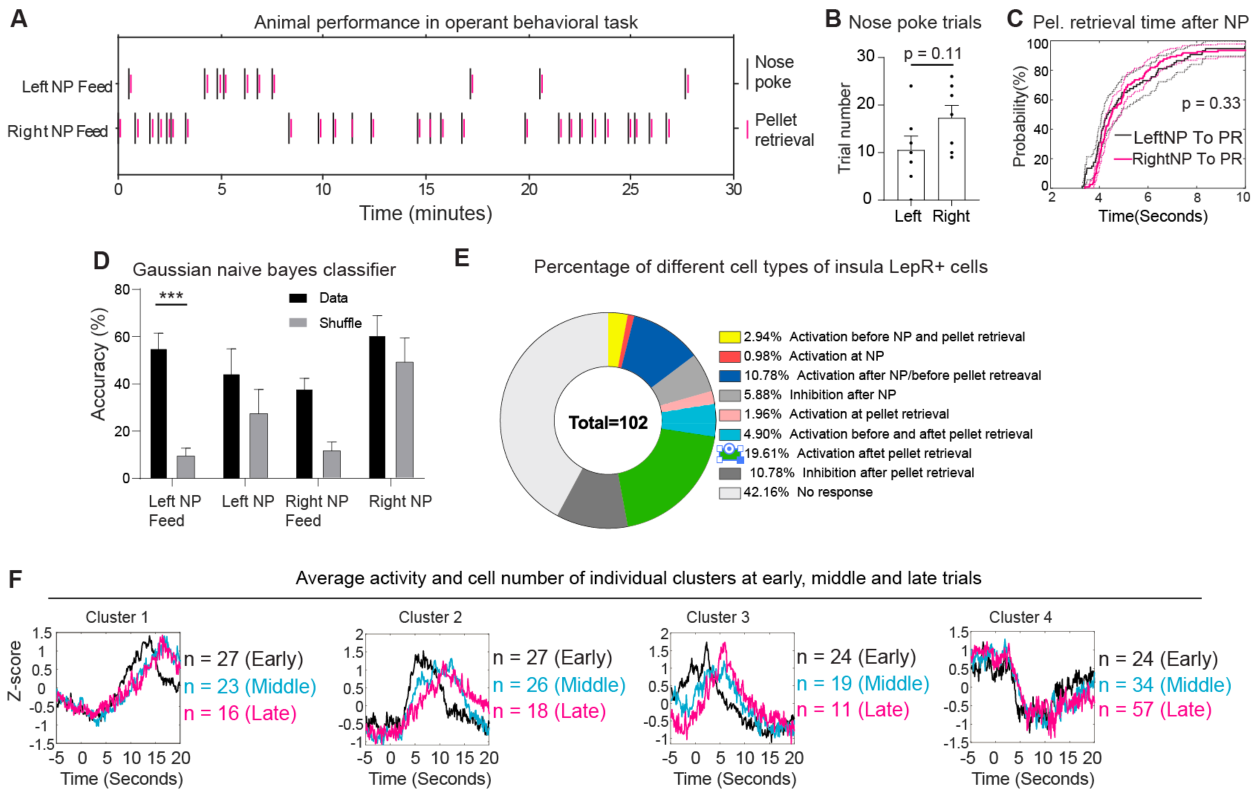


**Figure S4.** Neural dynamics of INS$^{LepR}$ cells in response to operant feeding behavior, related to **Figure 3**.

(A) Representative animal performance in operant behavioral task.
(B) Trial numbers of left nose poke and right nose poke in individual mice were not significantly different (n = 6 mice, left nose poke; n = 6, right nose poke, two-tailed paired t test, $p = 0.1112$).
(C) Cumulative distribution probability of pellet retrieval time after left nose poke (black) and right nose poke (magenta) did not differ (n = 74 from 6 mice, left nose poke; n = 121 from 6 mice, right nose poke, two-sample Kolmogorov-Smirnov test, $p = 0.329$).
(D) Gaussian naïve bayes classifier trained by the population of LepR+ cell activity can be used to predict a portion of the animal behavior (n = 6 data, n = 6 shuffle, data vs. shuffle, two-way ANOVA followed by Tukey post-hoc test, $F_{(1,40)} = 21.56$, **** $p < 0.0001$; decoding accuracy for Left NP Feed, *** $p < 0.0006$).
(E) Percentage of specific insula LepR+ cell responses during operant feeding.
(F) Average activity and number of cells in individual clusters at early, middle and late trials. Early: black; Middle: cyan; Late: magenta.

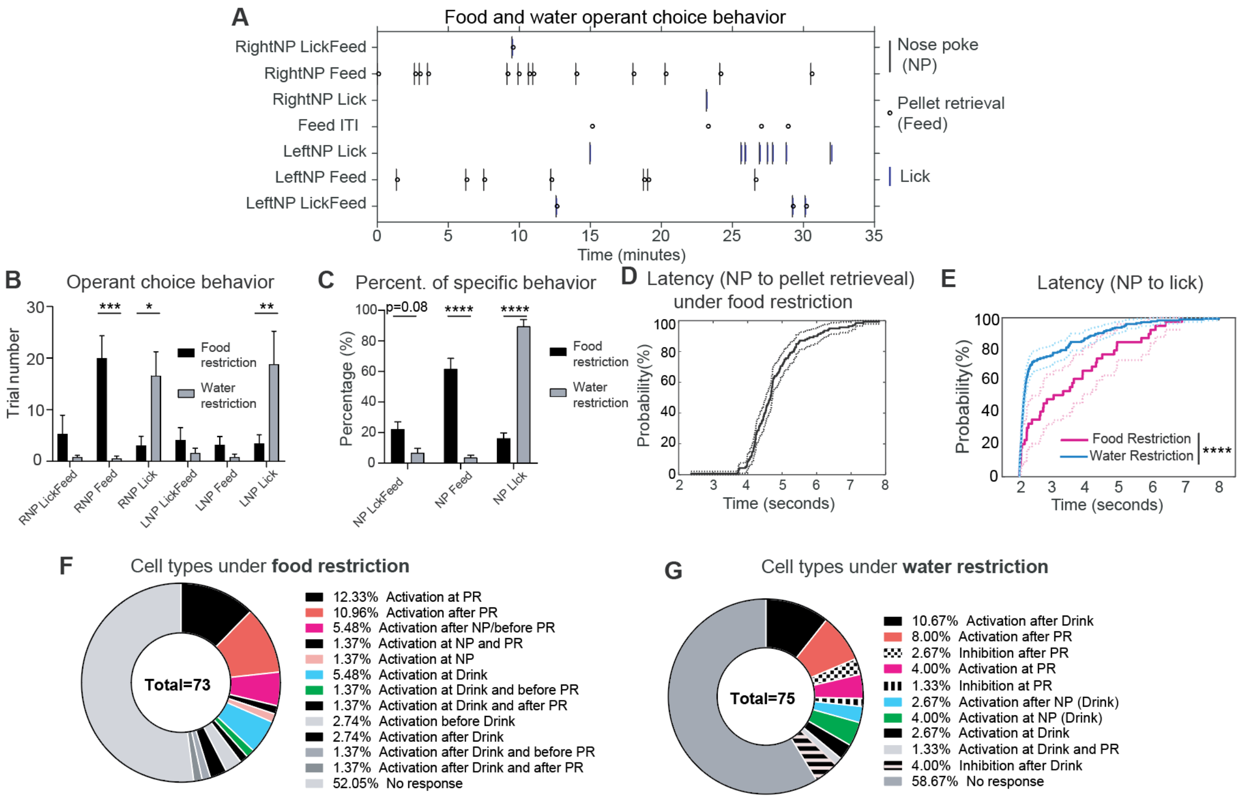


**Figure S5.** Neural dynamics of $INS^{LepR}$ cells in operant choice behavior, related to **Figure 4**.

(A) Representative animal performance in operant behavioral task. ITI: intertrial interval
(B) Number of trials for distinct behavioral choices: right/left nose poke, followed by Lick+Feed/Feed/Lick in individual mice under food (black) or water (grey) restriction (n = 6, Food restriction; n = 5, Water restriction, two-way ANOVA followed by Tukey post-hoc test, $F_{(1,54)} = 1.03 \times 10^{-5}$, $p = 0.9975$; RNP Feed, *** $p = 0.0002$; RNP Lick, * $p = 0.0135$; LNP Lick, ** $p = 0.004$).
(C) Percentage of distinct behavioral choices in individual mice under food (black) or water (grey) restriction (n = 6, Food restriction; n = 5, Water restriction, two-way ANOVA followed by Tukey post-hoc test, $F_{(1,27)} = 4.085 \times 10^{-14}$, $p > 0.9999$; NP LickFeed, $p = 0.0771$; NP Feed, **** $p < 0.0001$; NP Lick, **** $p < 0.0001$).
(D) Cumulative distribution probability of pellet retrieval time after nose poke (latency) under food restriction.
(E) Cumulative distribution probability of lick time after left nose poke (latency) under food (magenta) and water (blue) restriction (n = 39 from 6 mice, food restriction; n = 177 from 6 mice, water restriction, two-sample Kolmogorov-Smirnov test, **** $p = 7.1144 \times 10^{-5}$).
(F-G) Percentage of specific insula LepR+ cell responses during operant feeding and drinking under food (**F**) or water (**G**) restriction.

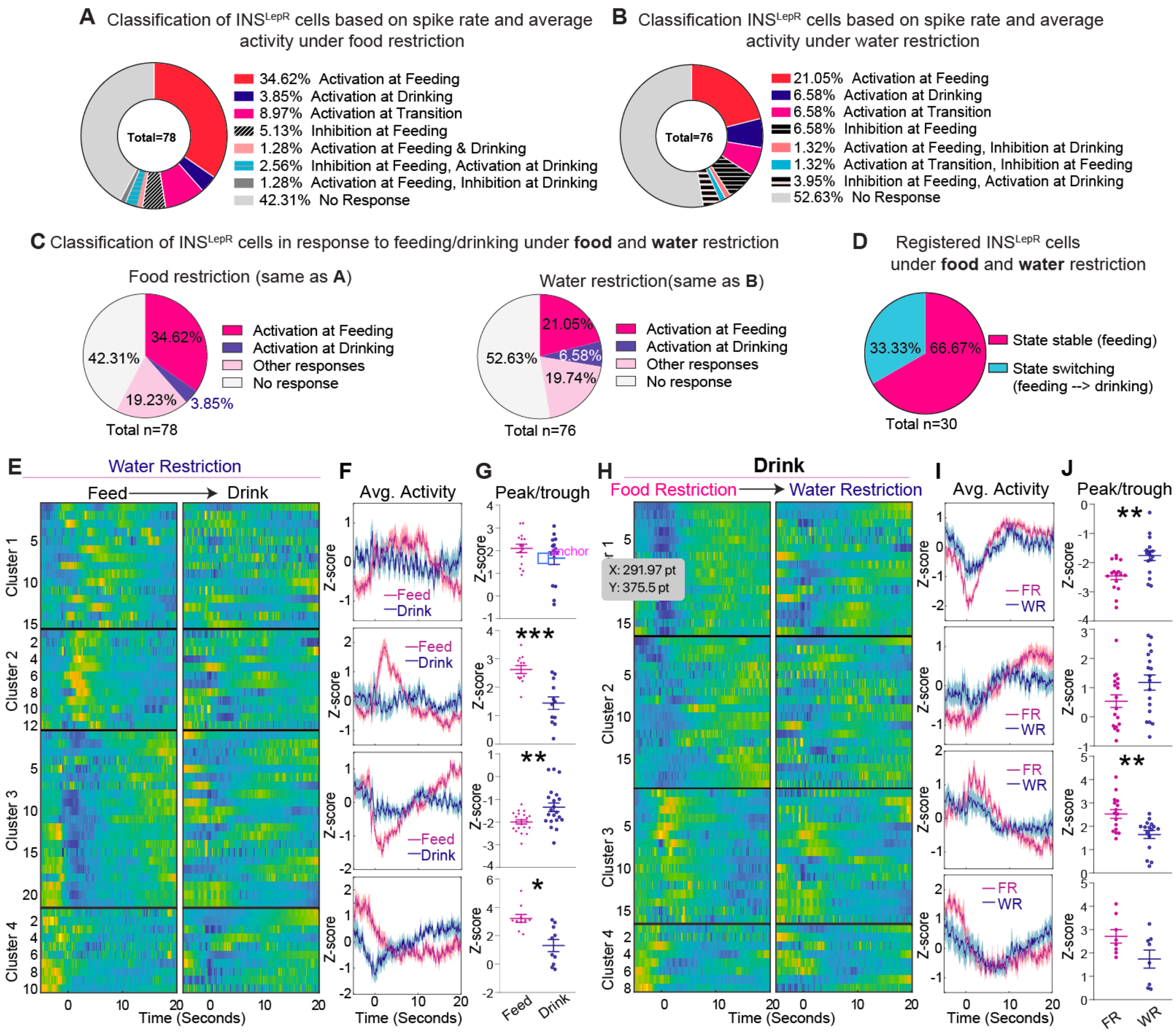


**Figure S6.** Classification of INS$^{LepR}$ cells in response to free feeding and drinking behavior, related to **Figure 5**.

(A-B) Classification of INS$^{LepR}$ cell response types during free feeding and drinking based on spike rate and average activity under food (**A**) and water (**B**) restriction.
(C) Cumulative classification of INS$^{LepR}$ cell responses to free feeding under food and water restriction, same data as **A** and **B**.
(D) Percentage (%) of registered INS$^{LepR}$ cells under food and water restriction, which show either a stable response to feeding (pink) or shifting response from feeding to drinking (blue).
(E) Classification of INS$^{LepR}$ cells aligned to feeding or drinking, from 6 water-restricted mice.
(F) Average (avg.) activity, in Z-score, of different clusters during water restriction in response to feeding (magenta) or drinking (blue). Activity is aligned at pellet retrieval or licking onset.

(G) Peak or trough activity, measured in Z-score, of individual cells in 4 clusters during water restriction, in response to feeding (magenta) or drinking (blue). For **G**, statistics is using two-tailed unpaired t test, cluster2, n = 12 feeding, n = 12 drinking, *** $p = 0.0002$; cluster3, n = 21 feeding, n = 21 drinking, ** $p = 0.004$; cluster4, n = 10 feeding, n = 10 drinking, ** $p = 0.0014$.

(H-J) $INS^{LepR}$ cell responses to free drinking under food and water restriction (H). Activity is aligned at drinking onset. Same analysis as **F-G**. For **J**, statistics were calculated using two-tailed unpaired t test, cluster1, n = 16 food restriction, n = 16 water restriction, ** $p = 0.0021$; cluster3, n = 16 food restriction, n = 16 water restriction, ** $p = 0.0016$.

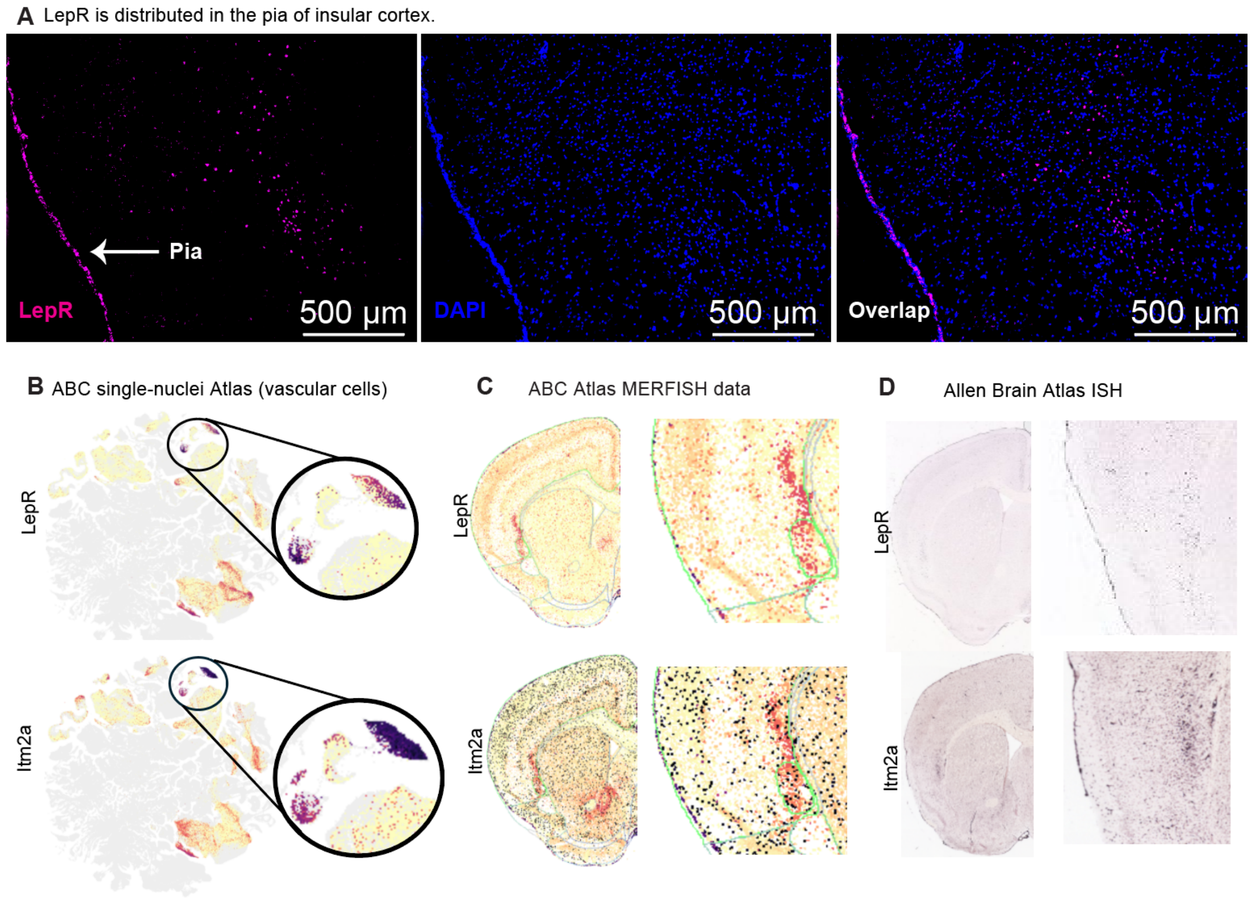


**Figure S7**. INS$^{LepR}$ cells co-express with markers of endothelial cells, related to **Figure 6**.

(A) LepR mRNA distribution in the pia of the insular cortex, as revealed by *in situ* hybridization.
(B) LepR+ and Itm2a+ expression level. Circular inset is magnified and shows clusters of vascular cells from the ABC single-nuclei sequencing atlas. Darker color indicates higher expression.
(C) LepR and Itm2a mRNA distribution reveals that both are expressed in deeper layers and claustrum, as well as along the pia. Darker color indicates higher expression. Data is taken from the ABC MERFISH Atlas
(D) LepR and Itm2a mRNA distribution shows similar expression to **C.** Data is taken from Allen Mouse Brain Atlas.

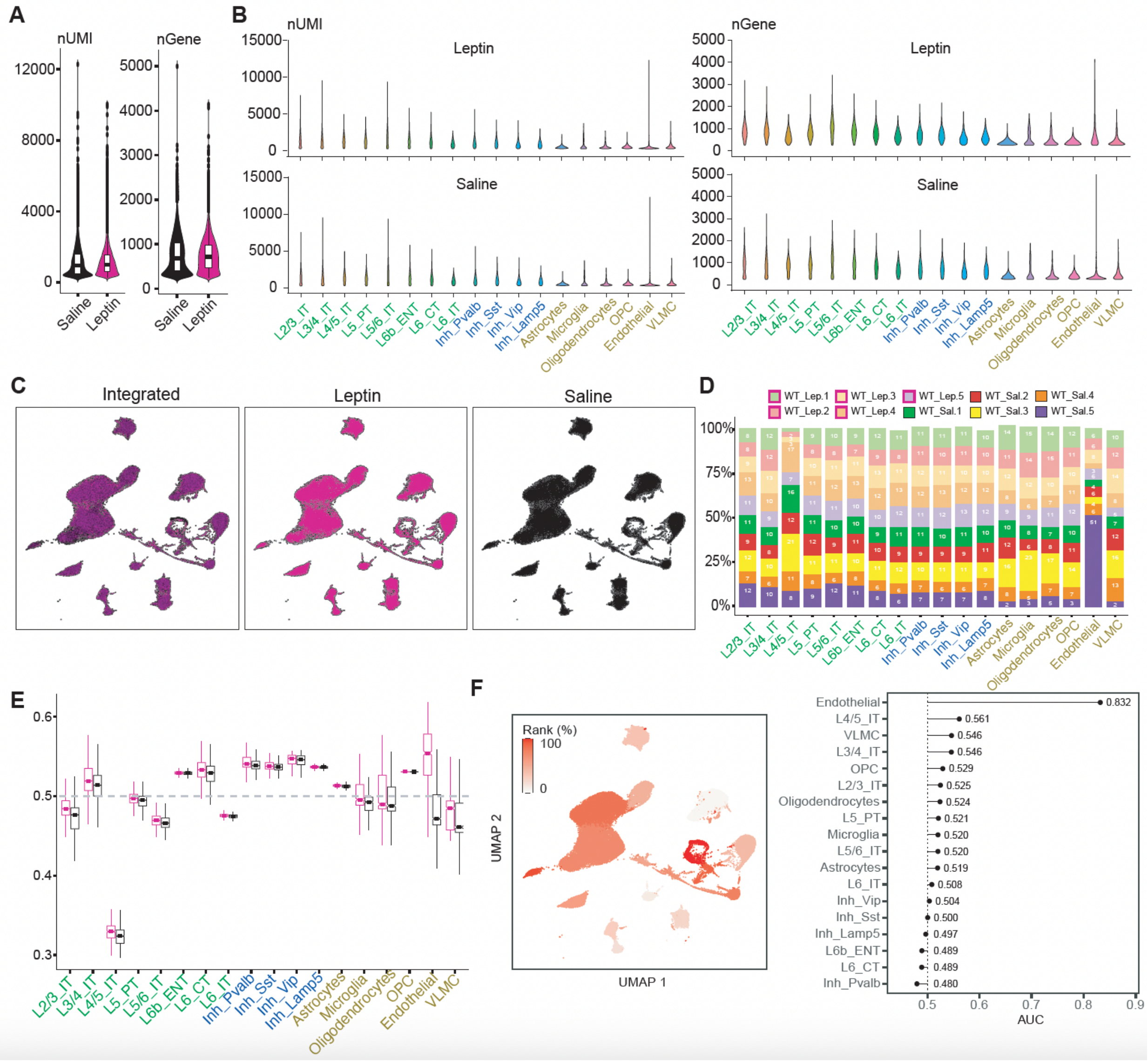


Figure S8. Genetic profiles of INS$^{LepR}$ cells, related to **Figure 6**.

(A-B) Distribution of UMI counts and number of genes in leptin treated and control mice (A) and per each cell type (B).
(C) UMAP showing the relative distribution of nuclei in leptin treated and control mice.
(D) Normalized proportion of each leptin treated and control mice samples for the cell type identified.
(E) Change in perturbation likelihood by leptin treatment across major cell classes calculated by MELD.
(F) AUGUR analysis of cell type prioritization in leptin treated compared with control mice. UMAP showing the AUGUR ranks (left). Lollipop chart depicting the AUGUR classifier score for each cell type (right).

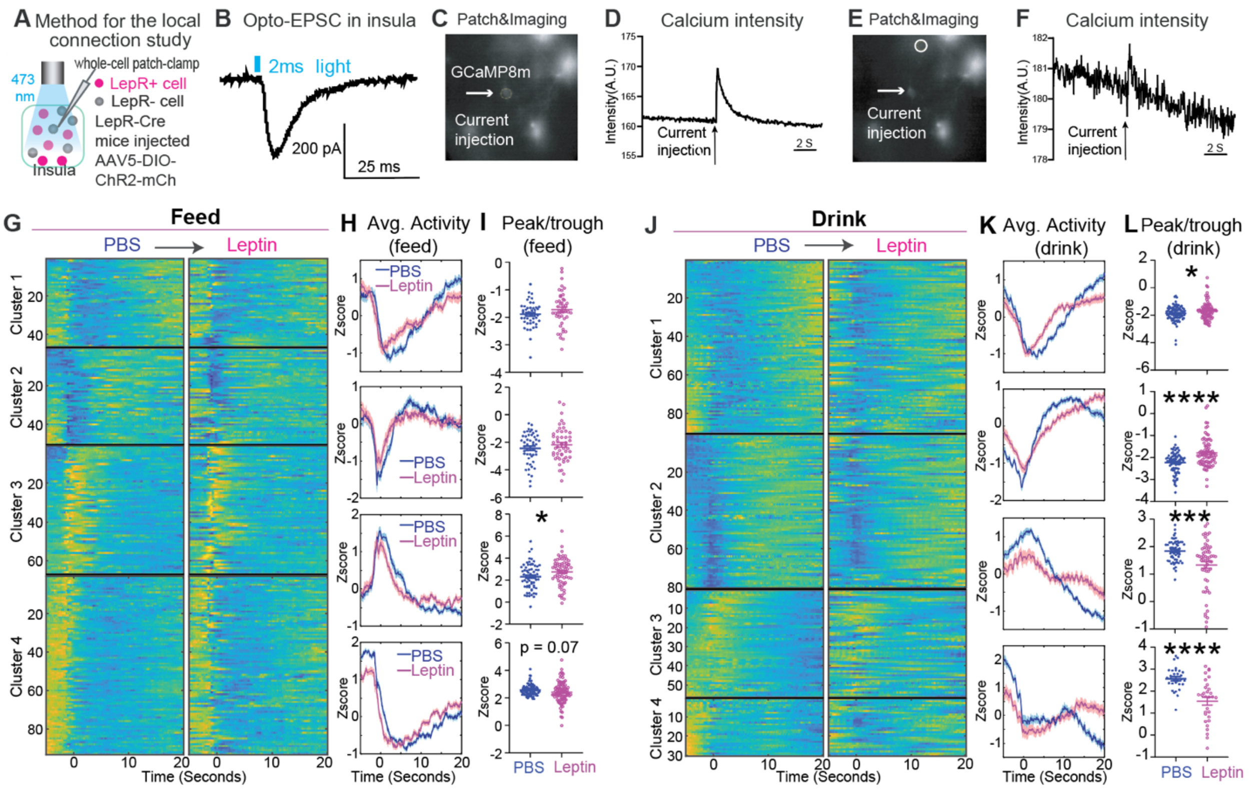

**Figure S9.** Functional connectivity of INS$^{LepR}$ cells and leptin shapes insula neural dynamics, related to **Figure 7**.

(A) Scheme for the method of studying functional connectivity by using whole-cell patch-clamp recording and optogenetic stimulation.
(B) Excitatory postsynaptic current (EPSC) in insula LepR- cell is induced by 2 ms of blue light on INS$^{LepR}$ cells expressing ChR2.
(C) Calcium imaging (bright white signal) in insula cells and stimulation of one patched single LepR+ cell (arrow) by current injection. Insula cells express GCaMP8m and LepR+ cells express mCherry.
(D) Calcium signals from circled cell in **C** increased after current injection.
(E-F) Calcium signal in a cell (white circle) above the patched cell increased after current injection.
(G) Classification of INS$^{LepR}$ cells aligned to feeding after PBS or leptin treatment from 6 mice under food restriction (G)
(H) Average activity, in Z-score of clusters in response to feeding under leptin (magenta) or PBS (blue) treatment. Activity is aligned at pellet retrieval.
(I) Peak or trough activity, in Z-score, of individual cells in 4 from **H**. For **I**, statistics is using two-tailed unpaired t test, cluster3, n = 67 PBS, n = 67 leptin, * $p$ = 0.0408.
(J) Classification of INS$^{LepR}$ cells aligned to drinking after PBS or leptin treatment, from 6 mice under food restriction.
(K) Average activity, in Z-score, of clusters in response to drinking under leptin (magenta) or PBS (blue) treatment. Activity is aligned at licking onsets.

(L) Peak or trough activity, in Z-score, of individual cells in 4 from **K**. For **L**, statistics is using two-tailed unpaired t test, cluster1, n = 90 PBS, n = 90 leptin, * $p = 0.0307$; cluster2, n = 80 PBS, n = 80 leptin, **** $p < 0.0001$; cluster3, n = 56 PBS, n = 56 leptin, *** $p = 0.0002$; cluster4, n = 30 PBS, n = 30 leptin, **** $p < 0.0001$.

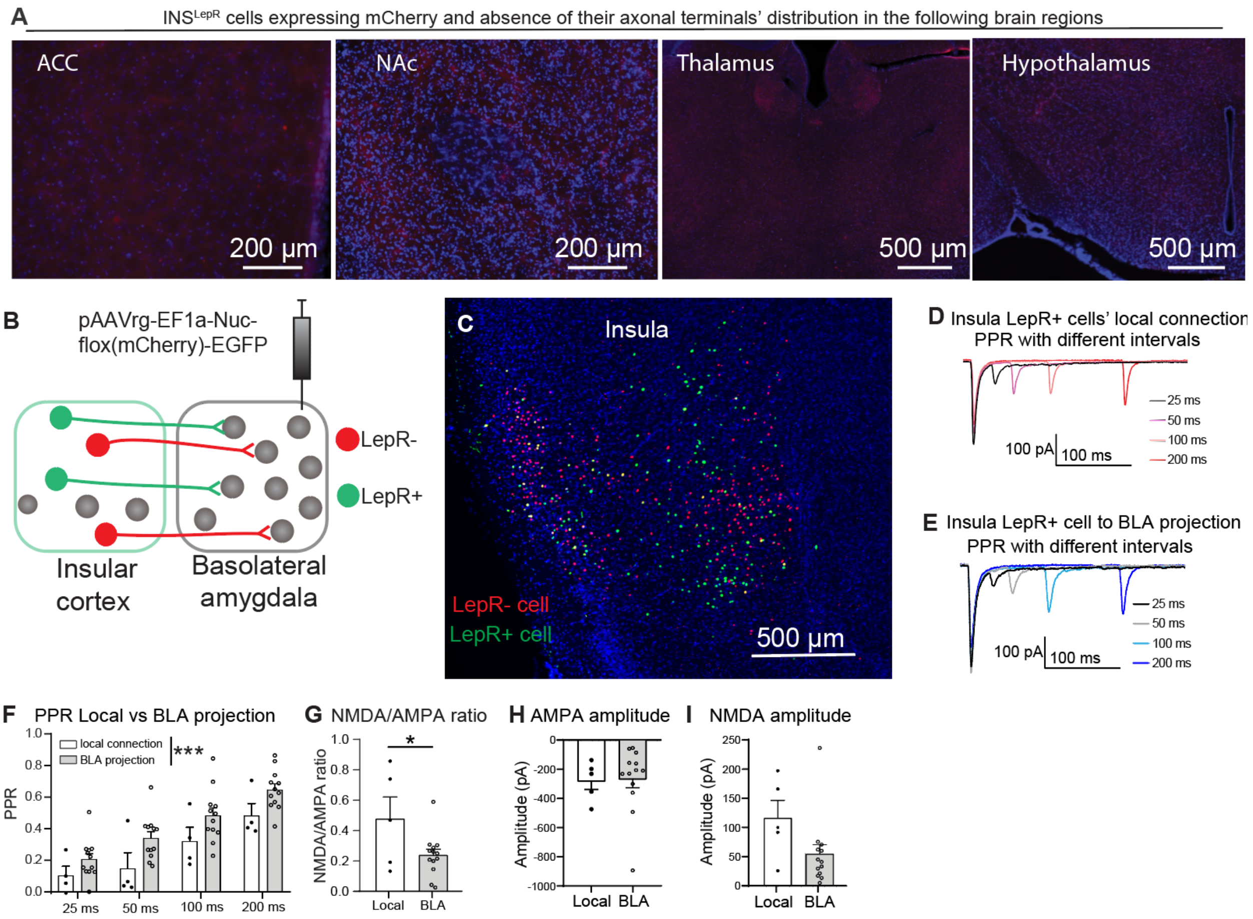


**Figure S10.** Functional connectivity of INS$^{LepR}$ cells from the insula to the basolateral amygdala, related to **Figure 7**.

(A) Absence of axonal terminals of the INS$^{LepR}$ cells expressing mCherry in the anterior cingulate cortex (ACC), nucleus accumbens (NAc), thalamus and hypothalamus.
(B) Scheme for injection of color-switching retrograde virus into the basolateral amygdala (BLA) of LepR-Cre mice. The virus expresses GFP in Cre+ neurons (e.g. LepR+) and mCherry in Cre- neurons (e.g. LepR-).
(C) LepR+ (green) and LepR- (red) cells in the insula which project to BLA, labeled by EGFP and mCherry, respectively.
(D) Paired-pulse ratio (PPR) of LepR- cells stimulated by LepR+ cells in the insula-claustrum.
(E) PPR of BLA cells stimulated by LepR+ cells in the insula.
(F) PPR of local LepR- cells and external BLA cells innervated by LepR+ cells in the insula (n = 4, local connection, n = 12 BLA projection, two-way ANOVA, $F(1,56) = 13.63$, *** $p = 0.0005$).
(G) NMDA/AMPA ratio of local LepR- cells and external BLA cells innervated by LepR+ cells in the insula-claustrum region (n = 5 local, n = 14 BLA, two-tailed unpaired t test, * $p = 0.0379$).
(H-I) AMPA (**H**) and NMDA (**I**) amplitudes of insula LepR- (local) and BLA cells innervated by INS$^{LepR}$ cells.
